# The ACF chromatin remodeling complex is essential for Polycomb repression

**DOI:** 10.1101/2021.06.29.450346

**Authors:** Elizabeth T. Wiles, Colleen C. Mumford, Kevin J. McNaught, Hideki Tanizawa, Eric U. Selker

## Abstract

Establishing and maintaining appropriate gene repression is critical for the health and development of multicellular organisms. Histone H3 lysine 27 (H3K27) methylation is a chromatin modification associated with repressed facultative heterochromatin, but the mechanism of this repression remains unclear. We used a forward genetic approach to identify genes involved in transcriptional silencing of H3K27-methylated chromatin in the filamentous fungus *Neurospora crassa*. We found that the *N. crassa* homologs of ISWI (NCU03875) and ACF (NCU00164) are required for repression of a subset of H3K27-methylated genes and that they form an ACF chromatin remodeling complex. This *N. crassa* ACF complex interacts with chromatin throughout the genome, yet association with facultative heterochromatin is specifically promoted by the H3K27 methyltransferase, SET-7. H3K27-methylated genes that are upregulated when *iswi* or *acf1* are deleted show a downstream shift of the +1 nucleosome, suggesting that proper nucleosome positioning is critical for repression of facultative heterochromatin. Our findings support a direct role for the ACF complex in Polycomb repression.

## INTRODUCTION

Polycomb repressive complex 2 (PRC2) methylates lysine 27 of histone H3 (H3K27), marking facultative heterochromatin (Muller et al. 2002; Margueron and Reinberg 2011). Facultative heterochromatin contains regions of the genome that must remain transcriptionally plastic in order to respond to developmental or environmental cues (Wiles and Selker 2017). Although H3K27 methylation has been established as a repressive chromatin modification, the mechanisms of repression are not fully understood (Margueron and Reinberg 2011; Ridenour et al. 2020). One model for repression involves Polycomb repressive complex 1 (PRC1) binding to the H3K27 methyl-mark to facilitate chromatin compaction by self-association (Grau et al. 2011; Cheutin and Cavalli 2018; Boyle et al. 2020). However, PRC1 is not present in all eukaryotes that bear H3K27 methylation-associated silencing, such as the filamentous fungus *Neurospora crassa* (Jamieson et al. 2013; Schuettengruber et al. 2017; Wiles et al. 2020), suggesting additional mechanisms of H3K27 methylation-associated repression.

In addition to histone modifications, nucleosome positioning may be critical for facultative heterochromatin function. The nucleosome, which is the fundamental unit of chromatin, consists of approximately 147 base pairs of DNA wrapped around an octamer of histone proteins (Kornberg 1974; Luger et al. 1997). Nucleosomes can be precisely positioned on DNA by ATP-dependent chromatin remodeling proteins to produce a chromatin landscape that modulates accessibility for DNA transactions, such as transcription (Lai and Pugh 2017). In particular, the precise positioning of the +1 nucleosome, the first nucleosome downstream of the transcription start site (TSS), is thought to be an important determinant of gene expression (Rhee and Pugh 2012; Nocetti and Whitehouse 2016). This dynamic nucleosome can occlude binding elements for transcriptional regulatory sites such as the TATA box (Kubik et al. 2018) and can serve as a barrier to RNA polymerase II (Weber et al. 2014).

In *Drosophila melanogaster*, the <u>A</u>TP-utilizing <u>c</u>hromatin assembly and remodeling factor (ACF) complex (Ito et al. 1997) has been indirectly linked to the repression of Polycomb targets (Fyodorov et al. 2004; Scacchetti et al. 2018). The ACF complex is comprised of the ATPase, Imitation Switch (ISWI), and the accessory subunit ACF1 (Ito et al. 1999). ACF is thought to act as a global nucleosome spacer and to contribute to repression genome-wide (Baldi et al. 2018; Scacchetti et al. 2018). Mutations in *Acf1* act as enhancers to *Polycomb* (*Pc*) mutations and disrupt nucleosome spacing in facultative heterochromatin (Fyodorov et al. 2004; Scacchetti et al. 2018).

In order to improve our understanding of the control and function of facultative heterochromatin, we used forward genetics to identify factors required for silencing H3K27-methylated genes in *N. crassa*. As described here, this identified *iswi* (also known as *crf4-1*) and *acf1* (also known as *crf4-2*; *Itc1* in *S. cerevisiae*). We show that these proteins interact to form an ACF complex in *N. crassa*. ACF is necessary for the repression of a subset of H3K27-methylated genes, but the derepression is not simply due to loss of H3K27 methylation. ACF interacts with chromatin targets throughout the genome, yet specific interactions with H3K27-methylated regions are partly dependent on SET-7, the H3K27 methyltransferase. Finally, we show that when members of ACF are absent, H3K27-methylated genes that become upregulated also display a specific downstream shift of the +1 nucleosome. Our findings support a model in which ACF remodels the chromatin landscape at H3K27-methylated regions of the genome to contribute to Polycomb silencing.

## RESULTS

### Forward genetic selection for genes required for Polycomb silencing identifies *iswi* and *acf1*

We previously designed and employed a forward genetic selection to identify novel genes required for silencing of H3K27-methylated genes (McNaught et al. 2020; Wiles et al. 2020). Briefly, the open reading frames of two genes (*NCU05173* and *NCU07152*) that require the H3K27 methyltransferase (SET-7) for repression were replaced with the hygromycin (*hph*) and nourseothricin (*nat-1*) resistance genes, respectively (Figure 1A). The resulting strain, which was sensitive to both antibiotics, was subjected to UV mutagenesis and grown on hygromycin- and nourseothricin-containing medium to select for drug resistant colonies. Drug resistant strains, which were in an Oak Ridge genetic background, were crossed to the polymorphic Mauriceville strain (Metzenberg et al. 1985). Spores from these crosses were germinated on hygromycin- and/or nourseothricin-containing media to select for the mutant segregants, and genomic DNA from the progeny was pooled for whole genome sequencing. Critical mutations were mapped by examining the ratio of Oak Ridge to Mauriceville single nucleotide polymorphisms (SNPs) genome-wide (Pomraning et al. 2011). Genetic variants were identified within the mapped regions using published tools (Danecek et al. 2011; Garrison and Marth 2012).

**Figure 1:**
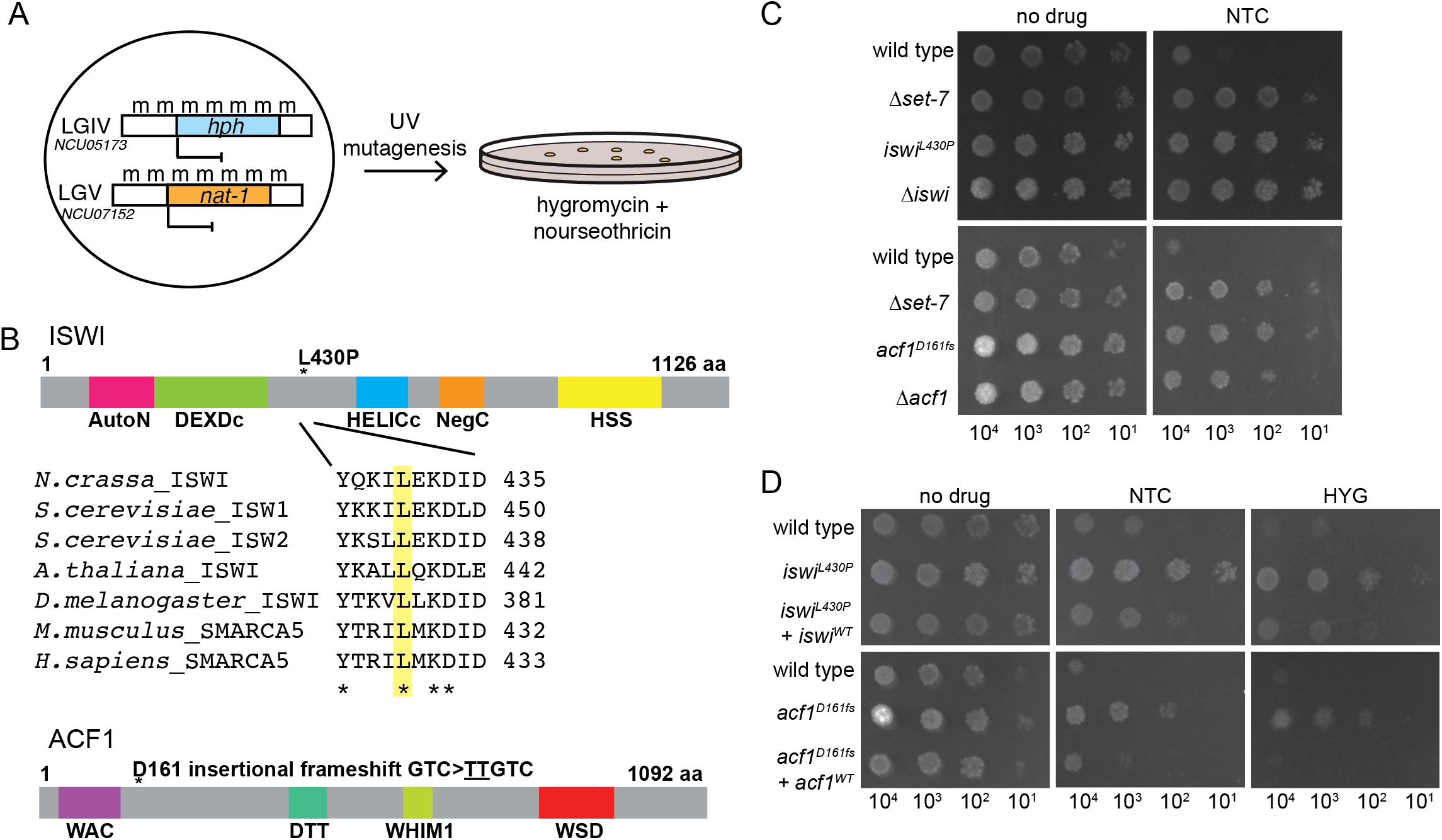
Forward genetics identifies ISWI complex members required for repression of H3K27 methylated genes. A, Selection scheme with reporter genes inserted at H3K27 methylation-marked loci to select for genes required for silencing. B, Schematic of protein domains in ISWI and ACF1 with the mutations identified in our selection (L430P and D161fs, respectively; marked with asterisks). The conserved nature of the mutated residue in ISWI is illustrated for the designated species. C, Serial dilution spot-test silencing assay for the indicated strains, which all contain *P_NCU07152_::nat-1* on media with or without nourseothricin (NTC). D, Serial dilution spot-test silencing assay for the indicated strains, which contain *P_NCU07152_::nat-1* and *P_NCU5173_::hph,* on media with or without nourseothricin (NTC) or hygromycin (HYG). Wild-type copies of each gene were inserted at the *his-3* locus for complementation. All spot tests were imaged after 48 h at 32°C and performed at least twice. The number of cells spotted is indicated beneath the images.

SNP mapping for one mutant identified a region on linkage group VI that contained essentially 100% Oak Ridge SNPs, indicating the likely position of the critical mutation (Figure 1 – figure supplement 1A). Within this region we found a point mutation (CTT -> CCT) in *iswi* (*NCU03875*) predicted to cause a leucine to proline substitution at a conserved position (L430P; Figure 1B). This same approach was used on a second strain to map and identify a two base pair insertion (GTC -> <u>TT</u>GTC) on linkage group III (Figure 1 – figure supplement 1B) that leads to a frameshift (D161fs) in *acf1* (*NCU00164*) (Figure1B).

To test if deletion of these two identified genes would also cause derepression of our H3K27 methylation mutant selection gene, we created strains with the *NCU07152::nat-1* replacement and either Δ*iswi* or Δ*acf1* alleles. Deletion of either *iswi* or *acf1* resulted in nourseothricin-resistance, equivalent to the original mutants identified in our selection (Figure 1C). In addition, we showed that introduction of an ectopic, wild-type copy of *iswi* or *acf1* into the corresponding original mutant strain largely restored silencing of both H3K27 methylation mutant selection genes (Figure 1D). We noticed that disruption of these two genes resulted in an early conidiation phenotype that was even evident in the spot tests (Figure 1C,D) but this was not accompanied by an increased linear growth rate. In fact, the Δ*iswi* strain showed a decreased growth rate relative to wild type; the Δ*acf1* strain grew comparably to wild type (Figure 1 – figure supplement 1C). Together, these data confirm that *iswi* and *acf1* are required to maintain the drug sensitivity of strains containing the H3K27 methylation mutant selection genes and are thus good candidates for genes involved in repression of H3K27-methylated chromatin.

### *iswi* and *acf1* are required for repression of a subset of H3K27-methylated genes

We performed mRNA-seq on Δ*iswi* and Δ*acf1* strains to determine if loss of these genes had specific effects on transcription within H3K27-methylated domains, or if they were general transcriptional regulators. We found that while the majority of gene expression changes observed upon loss of ISWI or ACF1 occurred outside of H3K27-methylated domains (Figure 2 – figure supplement 1A,B), genes marked by H3K27 methylation were significantly enriched in the upregulated gene sets for Δ*iswi* and Δ*acf1* strains (Figure 2A, B). Furthermore, nearly all (92%) of the H3K27-methylated genes that were upregulated in Δ*acf1* were also upregulated in Δ*iswi*, showing significant (P < 8.036E-54) overlap between these two gene sets (Figure 2C). This demonstrates that ISWI and ACF1 are not simply involved in the repression of the two H3K27-methylated genes we used in our initial selection (*NCU05173* and *NCU07152*), but also necessary for the repression of a large overlapping set of H3K27-methylated genes.

**Figure 2:**
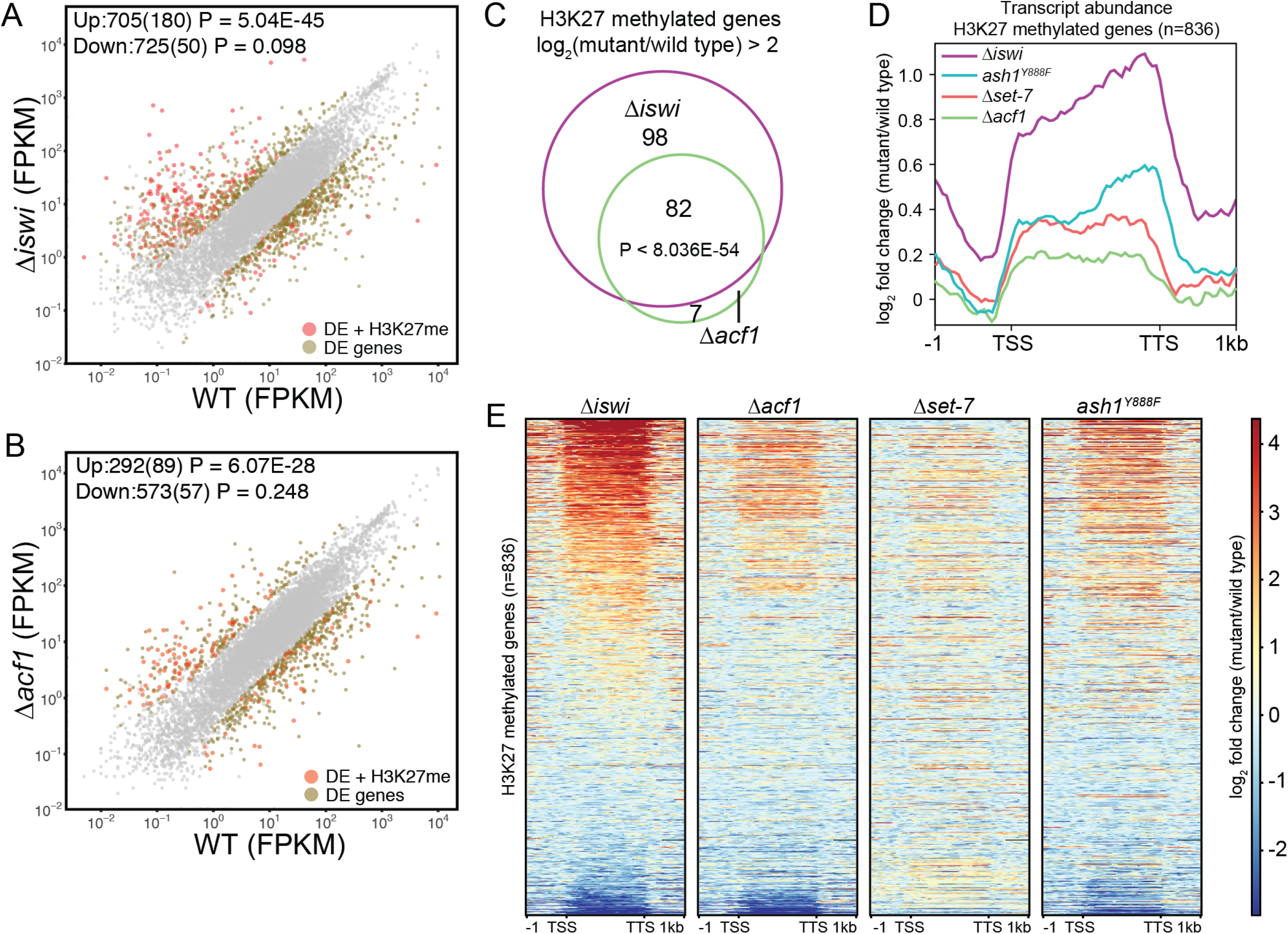
Loss of *iswi* or *acf1* leads to upregulation of a subset of SET-7-repressed genes. A and B, Expression level (FPKM) for each gene in the indicated mutant strain plotted against the expression level in wild type. Two biological replicates were used for each mutant and four biological replicates were used for wild type. Differentially expressed (DE) genes were defined using a significance cutoff of log_2_FC (fold change) > 2 for upregulated genes and log_2_FC < −2 for downregulated genes with a P value < 0.05. Gray dots indicate genes that are not considered differentially expressed. Upper left corner shows total number of significantly up- and downregulated genes with the number of H3K27-methylated genes in parentheses. Significance for enrichment of H3K27-methylated genes in each differentially expressed gene set was calculated by Fisher’s exact test. (FPKM - fragments per kilobase per million reads) C, Venn diagram showing overlap between H3K27-methylated genes that are upregulated (log_2_FC > 2; P value < 0.05) in Δ*iswi* and Δ*acf1* strains. Significant overlap (P < 8.036E-54) determined by hypergeometric probability test. D, Plots showing transcript abundance for all 836 H3K27-methylated genes for the indicated genotypes compared to wildtype. All genes are scaled to 2000 base pairs. E, Heat maps showing gene expression of all 836 H3K27-methylated genes for the indicated genotypes relative to wild type. All genes are scaled to 2000 base pairs. Genes are rank-ordered from most upregulated to most downregulated in Δ*iswi* and ordered the same in all heat maps. (TSS - transcription start site; TTS – transcription termination site)

Because we previously showed that the H3K27 methyltransferase SET-7 and the H3K36 methyltransferase ASH1 are involved in repression of H3K27-methylated genes, we compared the gene expression profiles from Δ*iswi* and Δ*acf1* strains to previously generated mRNA-seq data from Δ*set-7* or catalytic null ASH1 (*ash1^Y888F^*) strains (Figure 2D) (Jamieson et al. 2013; Bicocca et al. 2018). Interestingly, deletion of *iswi* resulted in a considerably greater increase in transcripts from H3K27-methylated genes than did either deletion of *set-7* or mutation of *ash1* (Figure 2D). Deletion of *acf1* strains showed a smaller increase in total mRNA abundance, even compared to the *set-7* or *ash1* mutants (Figure 2D). Increases in total transcript abundance can possibly result from a strong upregulation of few genes or a more modest upregulation of many genes. To investigate these possibilities, we created heatmaps that contained H3K27-methylated genes ordered from most upregulated to most repressed in Δ*iswi* strains and compared this rank-ordered list to gene expression changes in Δ*acf1*, Δ*set-7*, and *ash1^Y888F^* strains (Figure 2E). Consistent with our previous results, ISWI and ACF1 were found to be involved in the repression of a similar set of H3K27-methylated genes; however, this analysis revealed that deletion of *iswi* led to greater gene derepression. We also found that loss of ASH1 function derepresses a similar gene set as loss of either ISWI or ACF1, while fewer genes are upregulated in the Δ*set-7* strain. Closer inspection of overlap between gene sets shows that about half of Δ*set-7* upregulated genes are shared with upregulated genes in Δ*iswi* and Δ*acf1*, while the others are distinct (Figure 2 – figure supplement 1C). This furthers the notion that repression of H3K27-methylated genes is not simply a result of PRC2 activity.

### ISWI and ACF1 form a complex in *N. crassa*

Evidence from several organisms, most notably budding yeast and *Drosophila*, has implicated ISWI as the catalytic subunit of several chromatin remodeling complexes (Petty and Pillus 2013). To identify possible ISWI-containing protein complexes in *N. crassa*, we affinity-purified over-expressed 3xFLAG-ISWI from *N. crassa* cellular extracts. Immunopurified samples were digested down to peptides and analyzed by mass spectrometry (MS) to identify potential interacting proteins. We focused on proteins whose counts comprised greater than 0.4% of the total spectrum counts. We found that ISWI co-purifies with ACF1 as well as with CRF4-3 (NCU02684), a homolog of Ioc4 and member of the *Saccharomyces cerevisiae* Isw1b complex (Vary et al. 2003). Top hits from the MS results also included NCU00412 and NCU09388, proteins not known from *S. cerevisiae* (using NCBI BLASTP (Altschul et al. 1990)). Nevertheless, we considered that these could be members of ISWI complexes based on the high number of unique peptides detected and the presence of a WHIM domain in NCU00412 and a PHD domain in NCU09388 – domains that are present in *S. cerevisiae* Isw1 complex members, Ioc3 and Ioc2, respectively (Vary et al. 2003). NCU00412 and NCU09388 were also identified in a recent independent analysis of ISWI-interacting proteins in *N. crassa* and named ISWI accessory factor 1 and 2 (IAF-1 and IAF-2), respectively (Kamei et al. 2021). CRF4-3 was not previously identified as an ISWI-interacting partner, (Kamei et al. 2021) but for consistency, we will adopt the new nomenclature and refer to this protein as ISWI accessory factor 3 (IAF-3).

To confirm these interactions and to gain more information on the possible formation of ISWI-containing subcomplexes, we engineered a C-terminal HA tag at the endogenous locus of each of the four most prominent putative ISWI-interacting partners: ACF1, IAF-3, IAF-1 and IAF-2. These proteins were purified by immunoprecipitation and subjected to MS to identify interacting partners. Interactions between ISWI and all four proteins was confirmed, with each HA-tagged protein pull-down yielding high unique peptide counts for ISWI. Additional interactions, with lower unique peptide counts and lack of reciprocal pull-downs were also found (Figure 3A). These data suggest that ISWI forms multiple distinct protein complexes and, importantly, that ISWI and ACF1, two proteins identified in our selection for factors involved in repression of H3K27 methylated genes, interact. The ACF1-HA pull-down identified two histone fold proteins, NCU03073 (HFP-1) and NCU06623 (HFP-2)(Kamei et al. 2021), as interacting partners (Figure 3A). These proteins were also identified in our ISWI pull-downs, albeit below our threshold, and suggest the presence of an *N. crassa* CHRAC complex (Borkovich et al. 2004). The CHRAC complex is comprised of two histone fold proteins (DPB4 and DLS1 in *S. cerevisiae* and CHRAC14/16 in *D. melanogaster*) in addition to ISWI and ACF1 (Varga-Weisz et al. 1997; Corona et al. 2000; Iida and Araki 2004).

**Figure 3:**
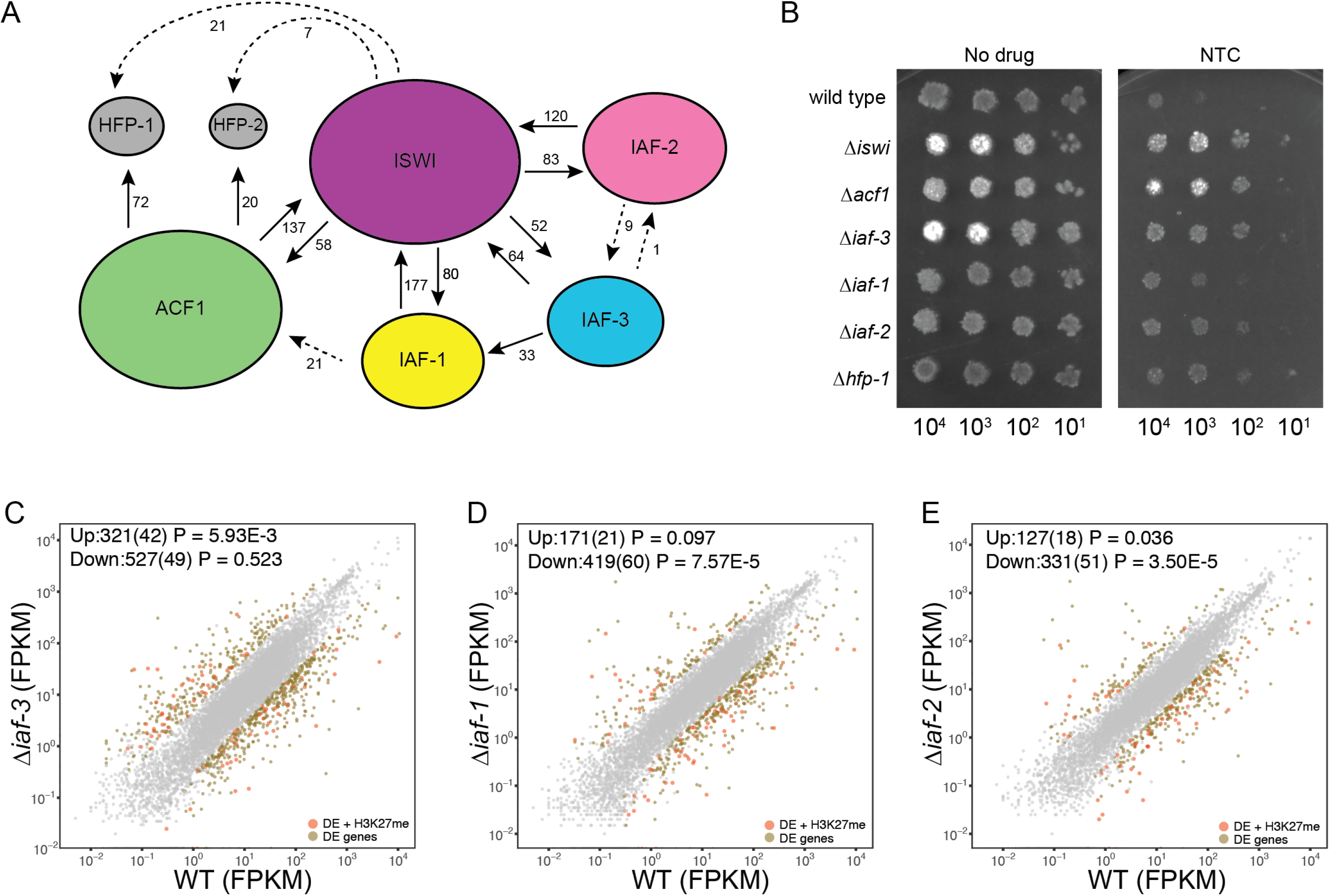
ISWI and ACF1 form a complex in *N. crassa.* A, Schematic representation of ISWI-interactions found by immunoprecipitation followed by mass spectrometry. Proteins (ISWI/NCU03875, ACF1/NCU00164, IAF-3/NCU02684, IAF-1/NCU00412, IAF-2/NCU09388, HFP-1/NCU03073, HFP-2/NCU06623) are depicted to scale. Arrows are drawn from the protein used as the “bait” to the protein partner identified, and unique peptide counts are indicated. Dotted arrows indicate the peptide count was below the 0.4% of total spectrum threshold. Proteins in gray (HFP-1 and HFP-2) were identified as interacting partners, but were not used as “bait.” B, Serial dilution spot-test silencing assay for the indicated strains on media with or without nourseothricin (NTC). All strains have *P_NCU07152_::nat-1*. The number of cells spotted is indicated beneath the images, which were generated after incubation for 48 h at 32°C. Spot test assays were repeated at least twice. C-E, Expression level (FPKM) for each gene in the indicated mutant strain plotted against the expression level in wild type. Two biological replicates were used for each mutant and four biological replicates were used for wild type. Differentially expressed (DE) genes were defined using a significance cutoff of log_2_FC (fold change) > 2 for upregulated genes and log_2_FC < −2 for downregulated genes with a P value < 0.05. Gray dots indicate genes that are not considered differentially expressed. Upper left corner shows total number of significantly up- and downregulated genes with the number of H3K27-methylated genes in parentheses. Significance for enrichment of H3K27-methylated genes in each differentially expressed gene set was calculated by Fisher’s exact test. (FPKM - fragments per kilobase per million reads)

To determine if any of the identified ISWI-interacting proteins, beyond ACF1, were involved in H3K27-methylated gene silencing, we first examined whether they were required for silencing the *NCU07152::nat-1* selection marker. As previously shown, deletion of *iswi* or *acf1* results in robust growth on nourseothricin indicating strong derepression of the *nat-1* gene. Deletion of *iaf-3* or *hfp-1* also derepressed the *nat-1* marker. Strains with deletion of *iaf-1* and *iaf-2* showed more modest growth on nourseothricin (Figure 3B). To determine the extent to which the other three ISWI-interacting partners may contribute to H3K27 methylation-mediated repression, we performed mRNA-seq on strains with deletions of *iaf-3*, *iaf-1*, or *iaf-2*. We found that H3K27-methylated genes were modestly enriched in the Δ*iaf-3* and Δ*iaf-2* upregulated gene sets (Figure 3C,E), but not enriched in the Δ*iaf-1* gene set (Figure 3D). Taken together, these findings suggest that ISWI forms multiple protein complexes in *N. crassa* and that the ACF complex makes the greatest contribution to repression of H3K27 methylated genes.

### *iswi* and *acf1* are required for H3K27 methylation at a subset of genes

To investigate if derepression of H3K27-methylated genes in Δ*iswi* and Δ*acf1* strains is due to loss of H3K27 methylation in these regions, we performed H3K27me2/3 ChIP-qPCR at the two H3K27 methyl-marked genes (*NCU05173* and *NCU07152*) used for our initial mutant selection. Both of these genes retained H3K27me2/3 (Figure 4A). To evaluate H3K27me2/3 distribution genome-wide, we performed ChIP-seq. We compared H3K27me2/3 at each gene and found, using two-fold cutoffs, that Δ*iswi* strains showed increased H3K27me2/3 for 218 genes and decreased H3K27me2/3 for 373 genes (Figure 4B). In contrast, Δ*acf1* strains showed only 3 genes with H3K27me2/3 gains and 80 genes with losses (Figure 4C). By comparing H3K27me2/3 changes between the two genotypes, we found that the vast majority of genes that changed H3K27me2/3 level in Δ*acf1* changed in the same direction in Δ*iswi* : 2 of 3 H3K27me2/3 gains and 77 of 80 H3K27me2/3 losses in Δ*acf1* were also changed in Δ*iswi*. By examining H3K27me2/3 genome-wide, we found that much of the distribution in Δ*iswi* and Δ*acf1* strains mirrored that of wild type (Figure 4D, Figure 4 – figure supplement 1A) while some regions differed from wild type and/or each other (Figure 4E, Figure 4 – figure supplement 1A). H3K27me2/3 losses were generally found at internal domains whereas gains were found at telomere-adjacent domains (Figure 4F,G). H3K27me2/3 ChIP-seq of strains with genes for other ISWI-interacting proteins deleted (Δ*iaf-3*, Δ*iaf-1*, or Δ*iaf-2*) showed only minor changes in H3K27me2/3 (Figure 4 – figure supplement 1B-D).

**Figure 4:**
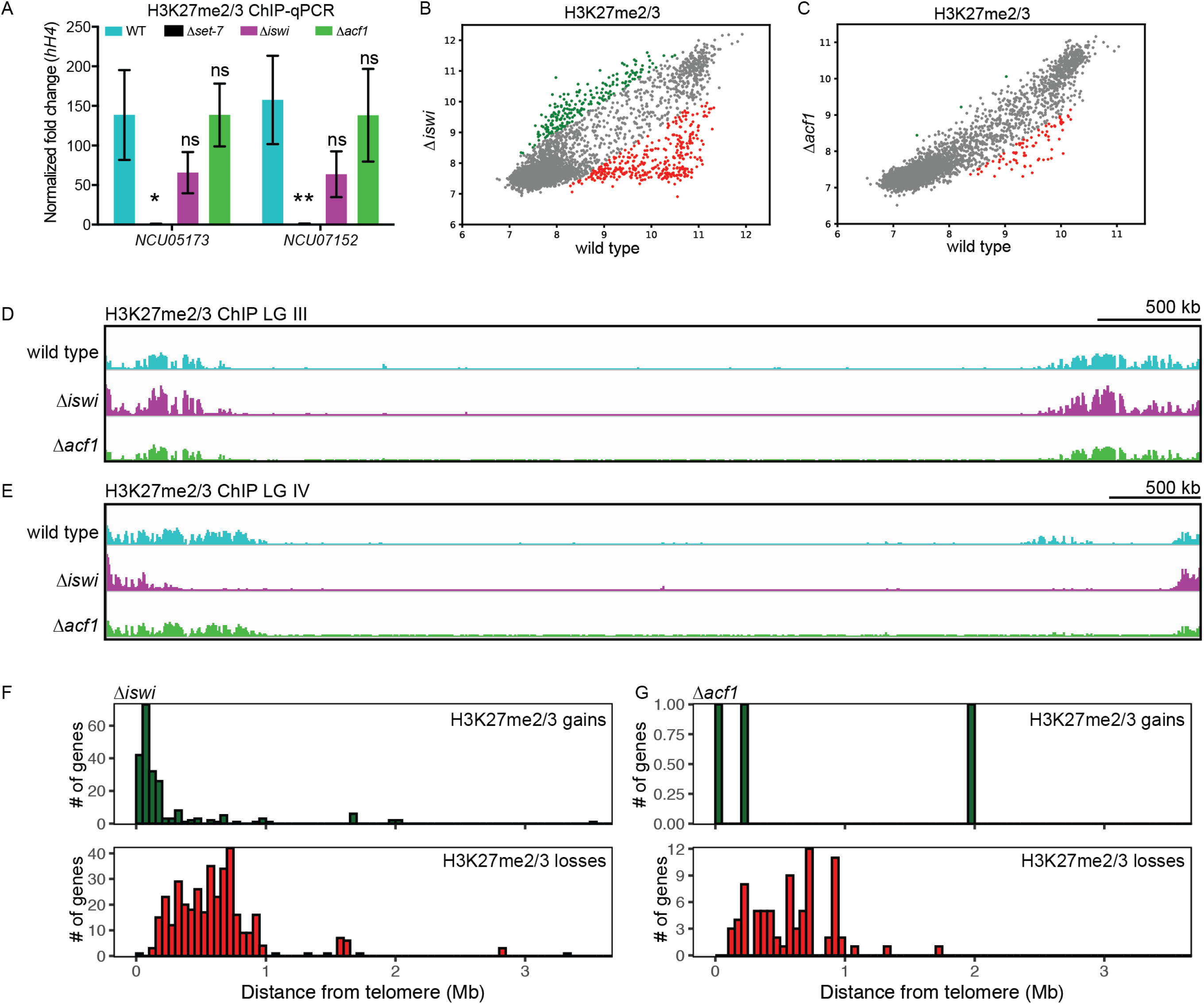
*iswi* and *acf1* are required for H3K27 methylation at a subset of genes. A, ChIP-qPCR data for H3K27me2/3 at the two genes used for the initial mutant selection (*NCU05173* and *NCU07152*) in the indicated strains. Filled bars represent the mean of technical triplicates and error bars show standard deviation (** for *P* < 0.01, * for *P* < 0.05, and ns for not significant; all relative to wild type by unpaired t-test). Data are from one representative experiment that was performed three times. B, Scatter plot showing correlation of H3K27me2/3 at genes in wild type and Δ*iswi* based on biological replicates of ChIP-seq data. Green points represent genes with increased H3K27me2/3 levels (at least 2-fold over wild type) and red points represent genes with decreased H3K27me2/3 levels (at least 2-fold relative to wild type) when *iswi* was deleted. C, Scatter plot as in (B) but for Δ*acf1*. D, ChIP-seq tracks showing average level of H3K27me2/3 from two biological replicates for the indicated strains on LG III. Y axis is 0-1000 RPKM for all tracks. E, Same as in (D) but for LG VI. F, Position of the H3K27me2/3 gains and losses in Δ*iswi* at each gene, as determined in (B), relative to the telomere. G, Same as in (F) but for Δ*acf1*.

Losses and gains of H3K27me2/3 alone do not account for all of the observed changes in gene expression (Figure 4 – figure supplement 1E,F). In Δ*iswi* strains, about one quarter of the genes that lost H3K27me2/3 were upregulated (100/373), while 16% (35/218) of genes that gained H3K27me2/3 were downregulated. In Δ*acf1* strains, about one third of genes (26/80) that lost H3K27me2/3 were also upregulated and one of the three genes that gained H3K27me2/3 was downregulated. These data show that loss of ISWI has a greater effect on H3K27 methylation than loss of ACF1, and that differences in H3K27 methylation do not account for all changes in gene expression in these deletion strains.

### SET-7 promotes ACF1 association with facultative heterochromatin

To identify chromatin targets of the *N. crassa* ACF complex, we fused the *E. coli* DNA adenine methyltransferase (van Steensel and Henikoff 2000) to the C-terminus of endogenous ACF1 and assayed adenine methylated DNA fragments by sequencing (DamID-seq) (Zhou 2012). We found that ACF1 localization is not restricted to one part of the genome, but rather appears to interact with chromatin indiscriminately genome-wide (Figure 5A, B). However, when *set-7* was deleted, eliminating H3K27 methylation, ACF1 localization to H3K27-methylated regions was reduced relative to wild type, suggesting that H3K27 methylation, or SET-7 presence, promotes ACF1 interactions specifically with these genomic regions (Figure 5A,B, Figure 5 – figure supplement 1A,B). These results were confirmed for two H3K27 methylation-marked regions (*NCU05173* and Tel VIIL) by Southern hybridizations with genomic DNA from DamID experiments. In contrast, deletion of *set-7* had no effect on ACF1-Dam localization at a euchromatic region (*his-3*) (Figure 5 – figure supplement 1A). When we compared ACF1-Dam localization to that of a nonspecific control (Dam only; referred to as Free-Dam), we found that both constructs localized to non-H3K27-methylated genes at similar levels, and this was independent of *set-7* presence (Figure 5C). In contrast, ACF1-Dam localized to H3K27-methylated genes more than Free-Dam and this increased localization was partially dependent on *set-7* (Figure 5D). These data suggest that ACF1 association with facultative heterochromatin is promoted by, but not fully dependent on, an intact PRC2 complex and/or H3K27 methylation.

**Figure 5:**
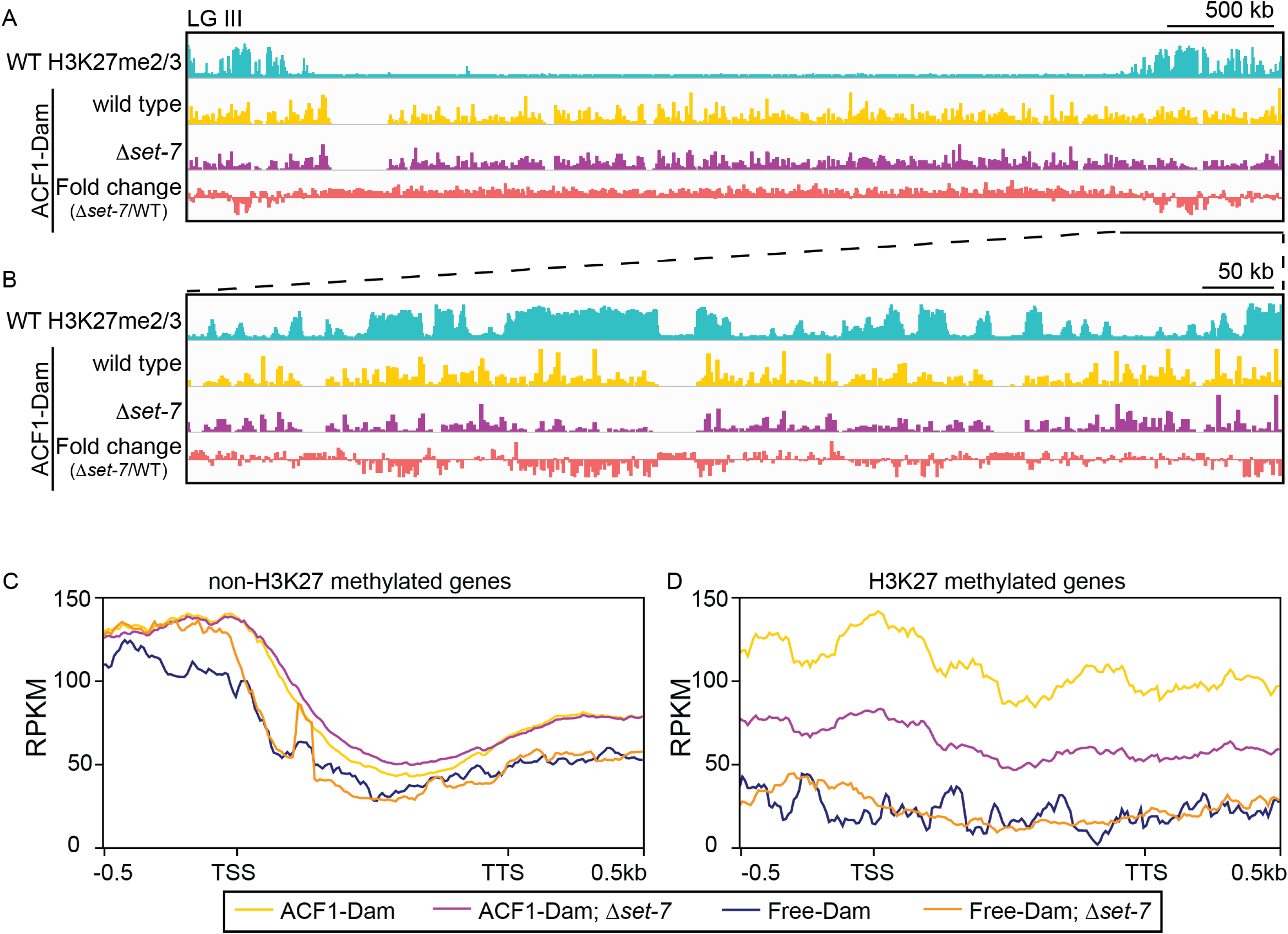
ACF1 localizes to H3K27me2/3-marked regions of the genome. A, Top track shows wild-type H3K27me2/3 levels based on ChIP-seq averaged from two biological replicates for an arbitrary chromosome (LG III). Y axis is 0-500 RPKM. Middle two tracks show DamID-seq average reads from two biological replicates for the indicated genotypes. Y axis is 0-500 RPKM. Bottom track compares the DamID-seq reads from Δ*set-7* strains to wild-type strains (shown above) displayed as the fold change between the two genotypes. Y axis is −3 to 3. B, Same as in (A) but showing an enlarged view of the right arm of LG III. Region shown is underlined in black in panel (A). C, Average enrichment based on DamID-seq for each non-H3K27-methylated gene, scaled to 1kb, +/-500 base pairs, is plotted for the indicated strains. All lines represent average reads from two biological replicates except for Free-Dam which is from only one. TSS – transcription start site; TTS – transcription termination site D, Same as in (C) but for H3K27-methylated genes.

### Loss of ACF has minor effects on nucleosome spacing

ACF-like complexes function differently in flies and yeast. In *D. melanogaster* ACF acts globally to space nucleosomes evenly (Baldi et al. 2018) whereas in *S. cerevisiae*, the analogous Isw2 complex specifically moves the +1 nucleosome in the 5’ direction, toward the nucleosome depleted region (NDR) (Whitehouse et al. 2007; Yen et al. 2012). To characterize nucleosome positioning in wild-type and mutant strains of *N. crassa, w*e performed MNase digestion followed by high throughput sequencing (MNase-seq). We first looked at nucleosome repeat length using the autocorrelation function (Braunschweig et al. 2009), which can analyze nucleosome positions independent of the transcription start site. When we looked genome-wide or considered only H3K27-methylated regions, we found only minor changes in nucleosome repeat length between wild-type and mutant strains (Δ*iswi*, Δ*acf1,* Δ*iaf-3*, Δ*iaf-1*, Δ*iaf-2* and Δ*set-7*) (Figure 6 – figure supplement 1A,B). This suggested that ISWI-containing complexes do not have major contributions to nucleosome spacing in *N. crassa* or there is redundancy among these proteins.

### Loss of ACF results in a downstream shift of the +1 nucleosome and transcriptional upregulation at a subset of H3K27-methylated genes

We next considered that the *N. crassa* ACF complex may function more like the *S. cerevisiae* Isw2 complex. For this analysis we looked at nucleosome positions in the promoter region of genes that had regular nucleosome arrays (defined as spectral density (SD) (Baldi et al. 2018) score > 2; n=7753) in at least one strain (wild type, Δ*iswi*, Δ*acf1,* Δ*iaf-3*, Δ*iaf-1*, Δ*iaf-2* or Δ*set-7*). We found that when all SD genes were considered, deletion of *iswi* or *acf1* was more likely to result in a downstream shift (>30 bp) of the +1 nucleosome than when *iaf-3*, *iaf-1*, *iaf-2* or *set-7* were deleted (Figure 6 – figure supplement 2A). This trend held when only H3K27-methylated SD genes (n=358) were considered (Figure 6A). Importantly, a significant portion of the H3K27-methylated genes with a shifted nucleosome is shared between *iswi* and *acf1* (P < 9.91E-13) (Figure 6B). These data suggest that ISWI and ACF1 may work in concert to position the +1 nucleosome at a subset of genes, including those in H3K27-methylated regions.

**Figure 6:**
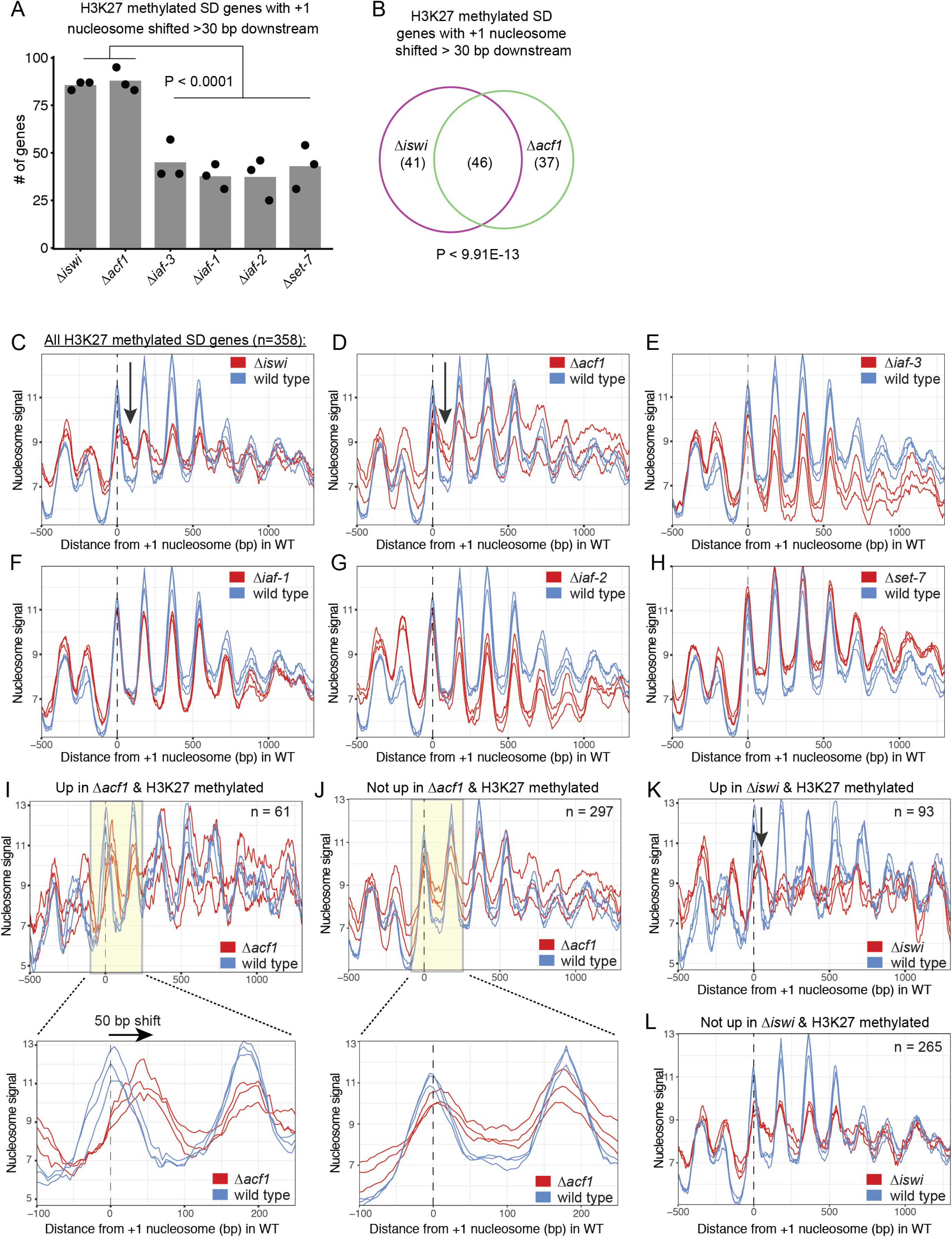
ISWI and ACF1 position the +1 nucleosome at H3K27-methylated, upregulated genes. A, Histogram of the number of H3K27 methylated SD genes (spectral density score for nucleosome order > 2; n=358) that have the +1 nucleosome shifted downstream >30 base pairs when compared to wild type in the indicated mutant strains. Each point represents biological replicate 1, biological replicate 2, or analysis of the merged replicates and filled bar is the average of all three values. P values were determined with an unpaired t-test. B, Venn diagram showing overlap of H3K27-methylated SD genes with a +1 nucleosome shifted downstream >30bp when *iswi* or *acf1* is deleted. P value was determined by hypergeometric probability test. C-H, Average nucleosome signal at all H3K27-methylated SD genes plotted from MNase-seq data for the indicated mutants and wild type. The three colored lines represent biological replicate 1, biological replicate 2, and the average of the replicates for the strains indicated in the key. Arrows in (C) and (D) indicate the shifted +1 nucleosome. I, Average nucleosome signal at SD genes that are upregulated (FDR < 0.05) and marked by H3K27 methylation in Δ*acf1* strains. The three colored lines represent biological replicate 1, biological replicate 2, and the average of the replicates. The boxed, shaded region is enlarged in the lower panel. J, Same as panel (I) but for H3K27-methylated SD genes that are not upregulated in Δ*acf1* strains. K, Average nucleosome signal at SD genes that are upregulated (FDR < 0.05) and marked by H3K27 methylation in Δ*iswi* strains. The three colored lines represent biological replicate 1, biological replicate 2, and the average of the replicates. Arrow indicates the shifted +1 nucleosome. L, Same as panel (K) but for H3K27-methylated SD genes that are not upregulated in Δ*iswi* strains.

Analysis of nucleosomes positions at all SD genes in wild-type and mutant strains (Δ*iswi*, Δ*acf1,* Δ*iaf-3*, Δ*iaf-1*, Δ*iaf-2* and Δ*set-7*) revealed some differences in occupancy at the −1 nucleosome but no global shift in nucleosome positions (Figure 6 – figure supplement 2B). Because such a genome-wide view can mask changes at specific targets and because the characteristic Isw2 5’ ‘pulling’ activity can only be appreciated when a subset of targets are examined (Yen et al. 2012; Ocampo et al. 2016; Donovan et al. 2021), we sought to limit our analysis to genes that might be targets of the ACF complex. Our inability to ChIP ACF1 and limitations of DamID-seq precluded a strict analysis of direct ACF1 targets but considering that our data support a functional role in transcriptional repression at H3K27-methylated genes, we restricted our analysis to these regions (H3K27-methylated SD genes; n=358). The MNase signal plots revealed that the +1 nucleosome shifted downstream in the absence of *iswi* or *acf1* (Figure 6C, D); in contrast, no shift was seen when other ISWI-interacting partners (Δ*iaf-3*, Δ*iaf-1*, Δ*iaf-2*) were deleted (Figure 6E-G). There was also no shift observed in Δ*set-7* strains when all H3K27-methylated SD genes were considered (Figure 6H), nor when we limited our analysis to H3K27-methylated SD genes that were upregulated upon deletion of *set-7* (Figure 6 – figure supplement 2C). These findings suggest that ISWI and ACF1 act to position the +1 nucleosome at a substantial subset of H3K27-methylated genes. Furthermore, SET-7 and hence H3K27 methylation are not required for nucleosome positioning by ISWI/ACF1 and transcriptional upregulation is not sufficient to shift nucleosomes.

To test if the downstream nucleosome shift at H3K27-methylated genes in Δ*iswi* or Δ*acf1* strains correlated with increased gene expression, we further focused our analysis to look at nucleosome positions in H3K27-methylated SD genes that were upregulated when *iswi* or *acf1* was deleted. We found that the +1 nucleosome shifted 50 bp downstream on average at H3K27-methylated SD genes that were upregulated (FDR < 0.05) in Δ*acf1* strains (Figure 6I) whereas no such shift was seen in the +1 nucleosome of H3K27-methylated SD genes that were not upregulated in Δ*acf1* strains (Figure 6J). Similarly, H3K27-methylated genes that were upregulated (FDR < 0.05) in Δ*iswi* display a more prominent downstream shift of the +1 nucleosome than those genes that were not upregulated (Figure 6K,L). Taken together, these data suggest that positioning of the +1 nucleosome by ISWI and ACF1 at a subset of H3K27-methylated genes contributes to transcriptional repression.

## DISCUSSION

### Control and function of facultative heterochromatin reflect the superimposition of a constellation of molecular mechanisms

Pioneering work on the Polycomb system of *Drosophila* revealed that methylation of lysine 27 of histone H3, catalyzed by Enhancer of Zeste in the PRC2 complex, is associated with, and important for, gene repression in facultative heterochromatin (Kassis et al. 2017). Although much has been learned, the importance of Polycomb repression in development of multi-cellular organisms has stymied progress towards a full understanding of its control and function. Moreover, there are indications of variable underlying mechanisms. For example, the PRC1 complex, which is widely regarded as central to Polycomb function in *Drosophila* and higher organisms, is less conserved than PRC2, and at least in some organisms, is absent (Schuettengruber et al. 2017). Similarly, Polycomb Response Elements (PREs), cis-acting DNA sequences controlling the distribution of H3K27me in *Drosophila*, do not appear to be universal (Kassis and Brown 2013). The complexity and importance of the Polycomb system in multicellular organisms led us to dissect the control and function of H3K27 methylation in the filamentous fungus *N. crassa*. We defined the PRC2 complex of Neurospora, demonstrated that it methylates H3K27 in roughly 7% of the genome, and is necessary for repression of scores of genes even though it is not essential in this organism (Jamieson et al. 2013; McNaught et al. 2020). Utilization of special genetic resources for Neurospora revealed that the organism has at least two distinct forms of H3K27 methylation (Jamieson et al. 2018), namely: 1. position-dependent, associated with telomere regions and characterized by involvement of the Neurospora p55 homolog (NPF) and the PRC2 accessory subunit (PAS) (McNaught et al. 2020), and 2. position-independent, which is found interstitially and does not depend on NPF or PAS (Jamieson et al. 2018). We’ve also previously shown that ASH1, an H3K36 methyltransferase, is critical for maintaining repression of many genes, including most of those in facultative heterochromatin (Bicocca et al. 2018).

The non-essential nature of H3K27 methylation, and the convenience of Neurospora for genetics and biochemistry, allowed us to design and implement a powerful selection for mutants defective in silencing genes in facultative heterochromatin. This unbiased scheme revealed both expected factors required for repression, including members of the PRC2 complex, and unanticipated players such as EPR-1 (Wiles et al. 2020) and the ACF complex reported here. A particularly interesting general finding is that repression is not simply due to a linear pathway of factors. While some factors cooperate to maintain repression, results of RNA-seq revealed considerable variation in the spheres of influence of the various factors. For example, loss of the H3K27 methyl-mark itself, or of the apparent H3K27 methyl-reader, EPR-1, each lead to derepression of somewhat different subsets of H3K27-methylated genes (Wiles et al. 2020) while loss of elements of the ACF remodeling machine, leads to loss of silencing of a larger set of H3K27 methyl-marked genes, even without loss of this characteristic mark of facultative heterochromatin. The overall picture that is emerging is cartooned in Figure 7 with more specifics discussed below.

**Figure 7:**
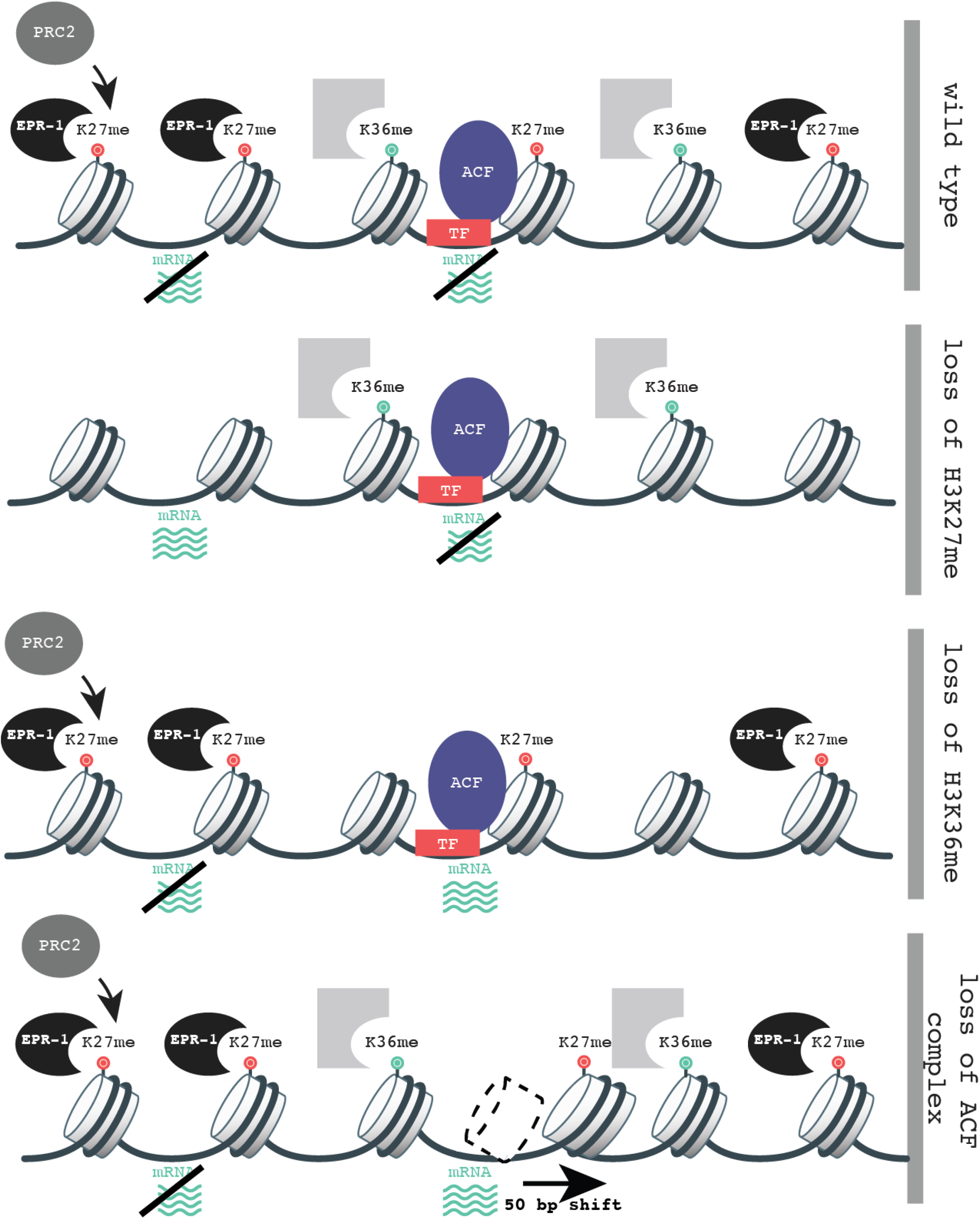
Multifaceted repression in facultative heterochromatin. A constellation of factors are responsible for maintaining silence of genes in regions marked by H3K27 methylation. Loss of this methyl-mark itself is sufficient to activate a fraction of genes, in part because of loss of the H3K27 methyl-specific factor EPR-1. Repression of many other genes, in H3K27 methylated domains and elsewhere, depend on both H3K36 methylation by ASH1 and both components of the ACF complex (ISWI and ACF). Gray partial square represents an unknown H3K36 methyl binding protein. TF represents unknown transcription factor(s) that could recruit/direct the ACF complex.

### The *Neurospora crassa* ACF complex is required for transcriptional repression at facultative heterochromatin

Our genetic identification of *iswi* and *acf1* as genes required for silencing H3K27 methyl-marked loci is consistent with a growing body of evidence that chromatin structure plays a major role in the transcriptional status of genes. Nucleosome-depleted regions (NDRs) are characteristic of transcriptionally active promoters and are thought to allow access of the transcriptional machinery (Lai and Pugh 2017). Conversely, nucleosomes can be positioned onto regulatory sequences in promoter regions by chromatin remodelers to cause repression (Whitehouse and Tsukiyama 2006; Whitehouse et al. 2007).

ACF-like complexes are conserved from budding yeast (Tsukiyama et al. 1999) to humans (LeRoy et al. 2000) but most of the biochemical studies of these complexes have been done with yeast and flies, which, curiously, revealed apparent functional discrepancies. In yeast, Isw2 (homologous to the ACF complex of *Drosophila*) acts in promoter regions where it binds to the +1 nucleosome and moves it in the 5’ direction toward the NDR (Whitehouse et al. 2007; Yen et al. 2012; Kubik et al. 2019). In contrast, ACF in *Drosophila* has been characterized as a nonspecific nucleosome spacing and assembly factor promoting global chromatin regularity (Baldi et al. 2018). The distinct modes of action of ACF-like chromatin remodelers in yeast and *Drosophila* warrant further study in other organisms. Our investigation of nucleosome positioning activities of ISWI and ACF1 in *N. crassa* revealed that these factors are required for positioning the +1 nucleosome at a subset of genes, including those marked by H3K27 methylation. Thus, *N. crassa* ACF complex seems to function more like the *S. cerevisiae* Isw2 than the *D. melanogaster* ACF. Although the detailed mechanism of recruitment and target selection for ACF-like complexes remains unclear, work in yeast implicates interactions of such complexes with transcription factors (Goldmark et al. 2000; Donovan et al. 2021). It was recently shown that the WAC domain of Itc1 in the Isw2 complex contains acidic residues required for binding to transcription factors and for nucleosome positioning at target promoters (Donovan et al. 2021). These residues (E33 and E40) are conserved in Neurospora (E32 and E39) but not in *Drosophila*, potentially accounting for the apparent less specific function of ACF in flies (Donovan et al. 2021). It will be of interest to determine if there are transcription factors that bind to facultative heterochromatin in *N. crassa* that mediate interactions with ACF1 to facilitate localization and direct activity of the ACF complex.

Our DamID-seq results are compatible with a “continuous sampling” model proposed for some ISWI chromatin remodelers (Erdel et al. 2010). In this model, the ACF complex transiently interacts with chromatin (Gelbart et al. 2005) throughout the nucleus in an autoinhibited conformation (Clapier and Cairns 2012; Ludwigsen et al. 2017) until some, still undefined, feature (Clapier and Cairns 2012; Hwang et al. 2014; Ludwigsen et al. 2017; Donovan et al. 2021) releases the autoinhibition and allows it to engage, activate, and move nucleosomes by hydrolyzing ATP. The transient nature of this chromatin interaction could account for our inability to confirm our ACF1 DamID findings with ChIP. Attempts to identify the targets of the homologous complex by ChIP have been also unsuccessful in *Drosophila* (Scacchetti et al. 2018).

### The *Neurospora crassa* ACF complex positions the +1 nucleosome in promoters of H3K27-methylated genes to mediate transcriptional repression

In theory, nucleosome movement could be either a cause or consequence of transcriptional activation. The fact that H3K27-methylated genes that are derepressed upon deletion of *set-7* did not show changes in the position of the +1 nucleosome suggests that changes in nucleosome position are not simply due to transcriptional activation. By extension, this supports the idea that transcriptional derepression of H3K27-methylated genes in *iswi* and *acf1* strains is a consequence of a misplaced +1 nucleosome. It is noteworthy that while Isw2-mediated repression is thought to occur by placement of the +1 nucleosome over important DNA regulatory elements, occluding transcriptional machinery and/or general regulatory factors (Whitehouse et al. 2007; Yen et al. 2012), full repression at some targets, such as the early meiotic genes, also requires histone deacetylase activity from Rpd3 (Goldmark et al. 2000; Fazzio et al. 2001). Clearly, the mechanism of repression by ACF, including the identification of additional players, perhaps including transcription factors, histone deacetylases, and other chromatin modifying factors, deserves further study.

### Conclusions

Despite differences in the modes of action of *S. cerevisiae* Isw2 and *Drosophila* ACF, their biological outcomes are the same – transcriptional repression (Goldmark et al. 2000; Fyodorov et al. 2004; Ocampo et al. 2016; Scacchetti et al. 2018). We found that nucleosome positioning by the *N. crassa* ACF complex also leads to transcriptional repression, particularly at H3K27-methylated regions of the genome, establishing the ACF complex as a player in transcriptional repression characteristic of facultative heterochromatin. It will be valuable to determine if interplay between Polycomb-mediated repression and ISWI chromatin remodelers holds in other organisms. Interestingly, ACF has been indirectly linked to Polycomb repression in flies (Scacchetti et al. 2018), and notably, ISWI components were also identified in a screen for factors required for Polycomb repression in mammalian cells (Nishioka et al. 2018), raising the possibility that the role of the ACF complex in Neurospora is general.

## MATERIALS AND METHODS

### Strains, media and growth conditions

All *N. crassa* strains were grown as previously described (Wiles et al. 2020) and are listed in Supplemental File 1. Technical replicates are defined as experimental repeats with the same strain. Biological replicates are defined as experiments performed using a different strain with the same genotype.

### Selection for mutants defective in Polycomb silencing

The selection was carried out as previously described (Wiles et al. 2020). Briefly, conidia from strain N6279 were mutagenized with UV radiation and subjected to selection with Hygromycin B or Nourseothricin. Resistant colonies were grown and crossed to strain N3756 to generate homokaryons.

### Whole genome sequencing, mapping and identification of mutants

Whole genome sequencing, SNP mapping and identification of mutants was performed as previously described (Wiles et al. 2020). Briefly, antibiotic-resistant, homokaryotic mutants were crossed to a genetically polymorphic Mauriceville strain and approximately 15-20 antibiotic-resistant progeny were pooled and prepared for whole genome sequencing using the Nextera kit (Illumina, FC-121-1030). Mapping of the critical mutations was performed as previously described (Hunter 2007; Pomraning et al. 2011). FreeBayes and VCFtools were used to identify novel genetic variants present in pooled mutant genomic DNA (Danecek et al. 2011; Garrison and Marth 2012). All whole genome sequencing data are available on NCBI Sequence Reads Archive (PRJNA714693).

### Immunoprecipitation followed by mass spectrometry

Strains N7810 (*his-3*::*P_ccg_*::3xFLAG-*iswi*), N7971 (endogenous *acf1*-HA), N8071 (endogenous *iaf-3*-HA), N7973 (endogenous *iaf-1*-HA) and N8075 (endogenous *iaf-2*-HA) were grown and protein extracted as previously described (McNaught et al. 2020) except that 500 mL cultures were used. Purification of 3xFLAG-tagged protein was performed as previously described (McNaught et al. 2020). For HA-tagged proteins, the same procedure was used except that 20 ug of α-HA antibody (MBL 180-3) was bound to 400 ul equilibrated Protein A agarose (Invitrogen, 15918014) by rotating at room temperature for 1h and washed 3x with extraction buffer and protein was eluted 3x with 300 ul of 1 mg/mL HA peptide (ThermoFisher, 26184) in 1x TBS. Samples were sent to and processed by the UC Davis Proteomics Core Facility for mass spectrometry and analysis.

### RNA isolation, RT-qPCR, and mRNA-seq

Total RNA was extracted from germinated conidia as previously described (Wiles et al. 2020), and used for mRNA-seq library preparation (Klocko et al. 2016). Sequencing was performed by the University of Oregon Genomics and Cell Characterization Core Facility.

### mRNA-seq data analysis

Sequence reads were aligned to the *Neurospora crassa* genome (OR74A) using STAR program (version 2.7.3a). Total aligned reads per *N. crassa* gene were calculated using RSEM software (version 1.3.1) and normalized using DESeq2 software (version 1.24.0). Batch effects were corrected using R package, limma (version 3.44.1). FDR < 0.05 and abs(log2 fold-change) > 2 were used as a threshold to identify significantly up- or downregulated genes. All sequencing files are available on the NCBI GEO database (GSE168277).

### Chromatin immunoprecipitation (ChIP), ChIP-qPCR, and ChIP-seq

H3K27me2/3 ChIP using α-H3K27me2/3 antibody (Active Motif, 39536), which recognizes di- or trimethylated H3K27, was performed as previously described (Wiles et al. 2020). The isolated DNA was used for qPCR (see Supplemental File 2 for primers) or prepared for sequencing (Wiles et al. 2020). Sequencing was performed by the University of Oregon Genomics and Cell Characterization Core Facility.

### ChIP-seq data analysis

Mapping, visualization, and analysis of ChIP-sequencing reads was performed as previously described (Wiles et al. 2020). All sequencing files are available on the NCBI GEO database (GSE168277).

### DamID Southern hybridization and sequencing

Southern hybridization was carried out as previously described (Miao et al. 2000) with probes generated by PCR amplification (see Supplemental File 3 for primers) from wild-type *N. crassa* genomic DNA (*NCU05173*, TelVIIL) or plasmid pBM61 (*his-3*). Genomic DNA was prepared for DamID-seq as previously described (Zhou 2012) with the modifications we have reported (Wiles et al. 2020). Sequencing was performed by the University of Oregon Genomics and Cell Characterization Core Facility. The “Free Dam” strain had an N-terminal NLS(SV40) and a C-terminal 3xFLAG tag and was expressed from the *his-3* locus.

### DamID-seq data analysis

DamID-seq mapping and analysis was done using the Galaxy public server (Afgan et al. 2018). The Barcode Splitter was used to filter for reads with a GATC at the 5’ end and these reads were mapped using Bowtie2 (Langmead and Salzberg 2012). Files for biological replicates were merged using MergeBam. Merged bam files were used as input for bamCoverage (RPKM, 50bp bins) to generate bigwig files for viewing on IGV and running bigwigCompare. The output from bamCoverage was used with computeMatrix to generate files to use for plotProfile and output graphs. All sequencing files are available on the NCBI GEO database (GSE168277).

### MNase digestion and sequencing

*N. crassa* cells were grown and digested with micrococcal nuclease as previously described (Laura E McKnight 2019) with the following modifications. MNase (Takara) concentration was optimized for each strain to yield ∼80-90% mononucleosomes (20 units for N3752, N3753, N7966, N8018, N7990, N7992, N7988, N7989; 60 units for N4718; and 80 units for N4730, N6170, N6171, N8016, N8017). All digestions were for 10 minutes at 37C, RNase (40 ug) treatment was for 1.5 hours at 42C and proteinase K (200 ug) treatment was for 1 hour at 65 C. 10 ug of gel-purified mononucleosome DNA was prepared for high-throughput sequencing using the NEBNext DNA Library Prep Master Mix Set for Illumina (NEB). Sequencing was performed by the University of Oregon Genomics and Cell Characterization Core Facility.

### MNase-seq data analysis

Paired-end sequence reads were aligned to the *Neurospora crassa* genome (OR74A) using Bowtie2 (version 2.3.3) with the option “-q -p 4 -X 250 --no-discordant --no-mixed --no-unal.” Paired-end alignment reads with maximum 250bp distance gap between them were used in subsequent analysis. This length corresponds to mononucleosomes. Only correctly aligned paired-end alignment reads were filtered using samtools (version 1.5) commands “samtools view –hf 0×2 input.bam | grep –v “XS:i:” Dyad Coverage was calculated using the scripts (03_PNA_SDE.R) (Baldi et al. 2018). All sequencing files are available on the NCBI GEO database (GSE168277).

### Spectral density (SD) estimation

The spectral density (SD) score corresponds to periods of 182bp and was calculated using the scripts (cov2spec.R) (Baldi et al. 2018). SD score was normalized as Z-score : (log2(SD score) – average) / standard deviation. Regions with the average Z-score threshold of 2 were defined as the domain with a regular nucleosome array.

### Autocorrelation function

The autocorrelation function (Braunschweig et al. 2009) was calculated for the dyad coverage vectors for the lag length of 1,000 bp. Nucleosome repeat lengths were obtained by linear regression of the first and second autocorrelation peak positions with zero intercept. The slope of the regression was defined as repeat length.

### Estimation of +1 nucleosome position

The average score of dyad coverage vector for every 182 bp using the region −100 bp to +1000 bp from TSS was calculated for each gene. The closest peak from TSS was defined as +1 nucleosome position.

### Data Availability

All RNA-seq, ChIP-seq, DamID-seq and MNase-seq data generated in this study have been submitted to the NCBI Gene Expression Omnibus (GEO; https://www.ncbi.nlm.nih.gov/geo/) under accession number GSE168277. All whole genome sequencing data haven been submitted to the NCBI Sequence Read Archive (SRA, https://www.ncbi.nlm.nih.gov/sra) under accession number PRJNA714693.

## Acknowledgments

We thank J. Lyle and R. Morse for help in genetic mapping of UV-generated mutants; V. Bicocca for performing initial experiments characterizing ISWI and for comments on the manuscript; and J. McKnight and L. McKnight for providing guidance and reagents for rapid MNase digestion. We also thank T. Bailey, D. Donovan, L. McKnight, and K. Noma for helpful comments on the manuscript. This work was funded by the National Institute of General Medical Sciences (GM127142 and GM093061 to E.U.S.), American Heart Association (14POST20450071 to E.T.W.), and K.J.M. was partially supported by the National Institutes of Health (HD007348).

## Author contributions

E.T.W, C.C.M., K.J.M, and E.U.S conceived and designed the experiments and analyzed the data. E.T.W, C.C.M. and K.J.M. performed the experiments. H.T. performed the bioinformatic analysis. All authors contributed to writing the manuscript.

---

**Figure 1 – figure supplement 1:**
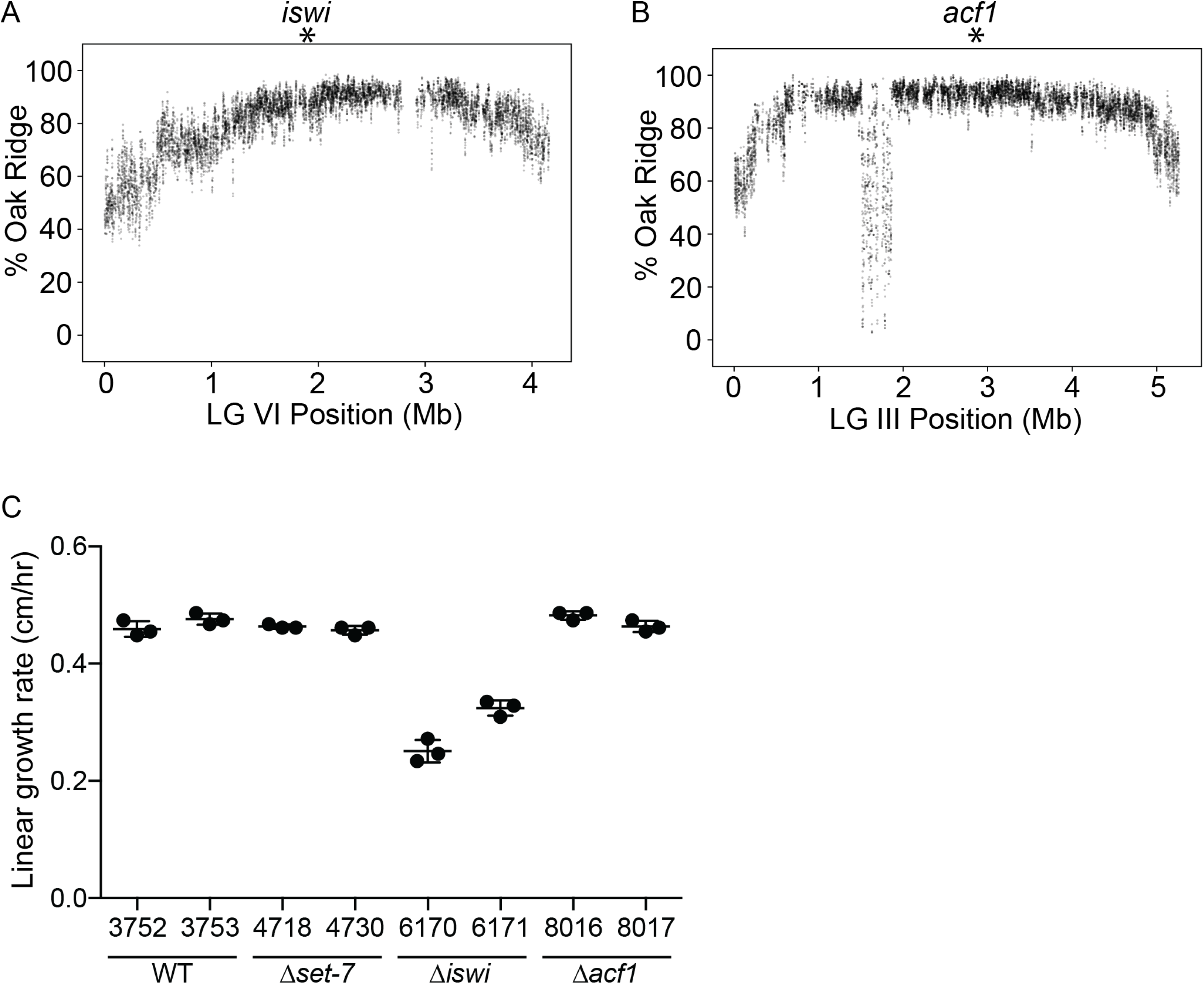
Genetic mapping and growth rate analysis of mutants identified in the selection. A, Whole genome sequencing of pooled mutant genomic DNA identified a region near the middle of LG VI (indicated by asterisk) that is enriched for Oak Ridge single nucleotide polymorphisms (SNPs) and contained a point mutation in *iswi*. Each point represents a running average (window size is 10 SNPs; step size is 1 SNP). B, Whole genome sequencing of pooled mutant genomic DNA identified a region near the middle of LG III (indicated by asterisk) that is enriched for Oak Ridge single nucleotide polymorphisms (SNPs) and contained a two base pair insertion in *acf1*. Each point represents a running average (window size is 10 SNPs; step size is 1 SNP). C, Linear growth rates as measured using race tubes for strains with the indicated genotypes. Points represent technical replicates.

**Figure 2 – figure supplement 1:**
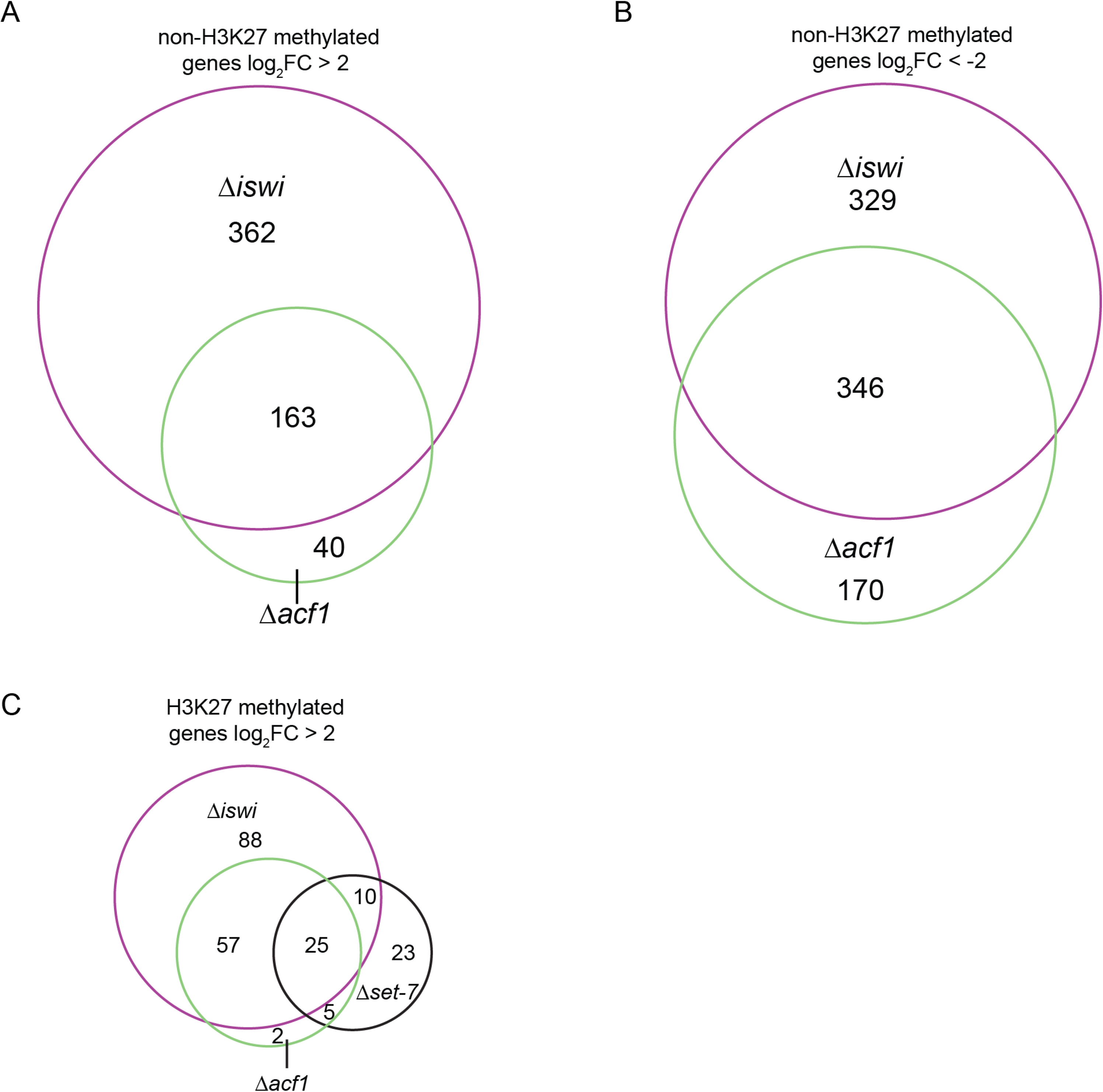
*iswi* and *acf1* are required for regulation of non-H3K27-methylated genes. A, Venn diagram showing the overlap of non-H3K27-methylated genes upregulated (log_2_FC > 2; P value < 0.05) for the indicated genotypes. B, Venn diagram showing the overlap of non-H3K27-methylated genes downregulated (log_2_FC < −2; P value < 0.05) for the indicated genotypes. Venn diagrams are not scaled relative to each other. C, Venn diagram showing the overlap of H3K27-methylated genes upregulated (log_2_FC > 2; P value < 0.05) for the indicated genotypes.

**Figure 4 – figure supplement 1:**
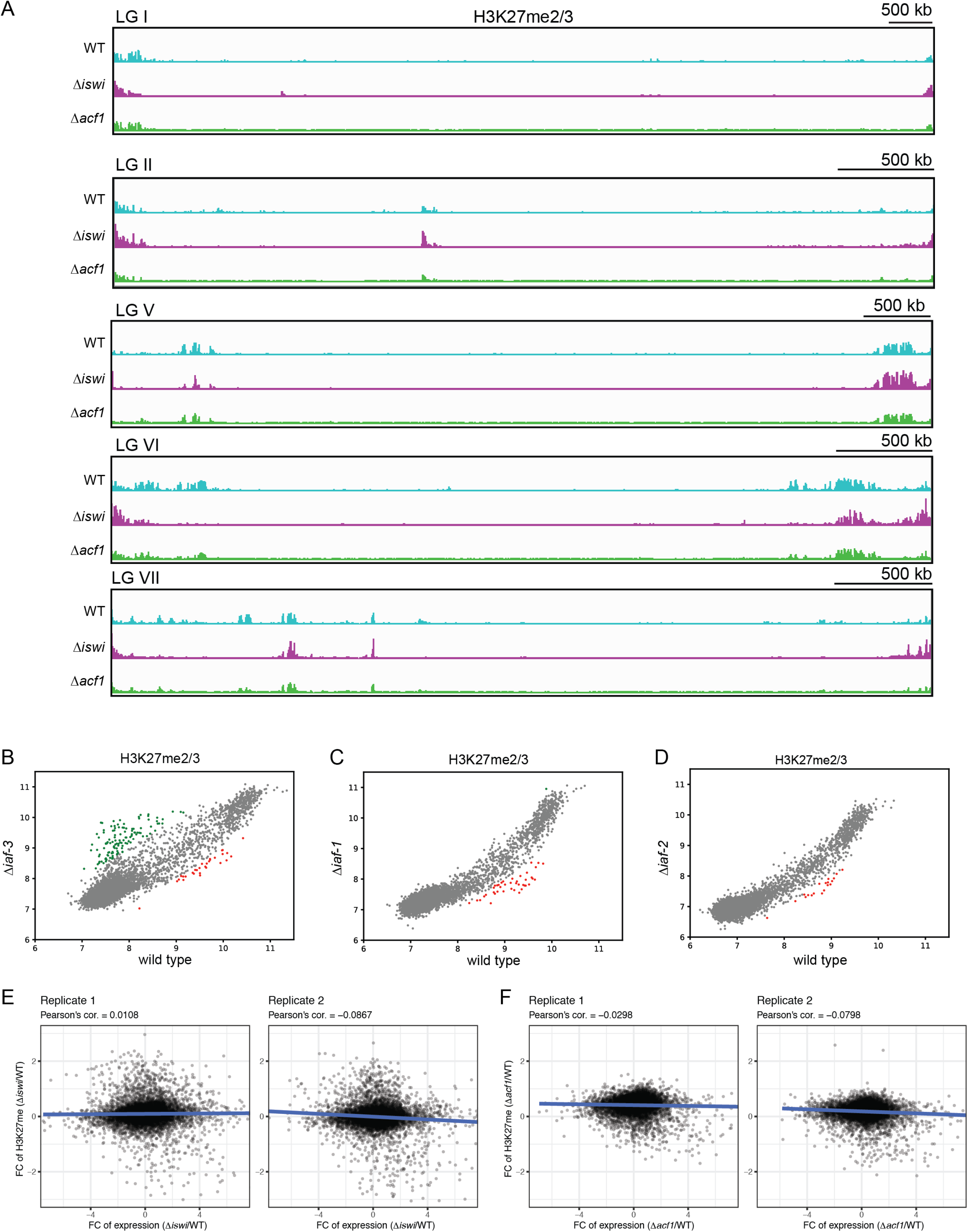
Loss of *iaf-3*, *iaf-1* and *iaf-2* results in minor changes in H3K27me2/3. A, ChIP-seq tracks showing average levels of H3K27me2/3 from two biological replicates for the indicated strains on the indicated chromosomes (linkage groups). Y axis is 0-1000 RPKM for all tracks. B, Scatter plot showing correlation of H3K27me2/3 at all genes in wild type and Δ*iaf-3* based on biological replicates of ChIP-seq data. Green points represent genes with increased H3K27me2/3 (at least 2-fold over wild type) and red points represent genes with decreased H3K27me2/3 levels (at least 2-fold relative to wild type) in Δ*iaf-3* strains. C, Same as in (B) but for Δ*iaf-1* strains D, Same as in (B) but for Δ*iaf-2* strains. E,F, Plots show correlation between fold change (FC) in H3K27 methylation and gene expression in Δ*iswi* and Δ*acf1* strains.

**Figure 5 – figure supplement 1:**
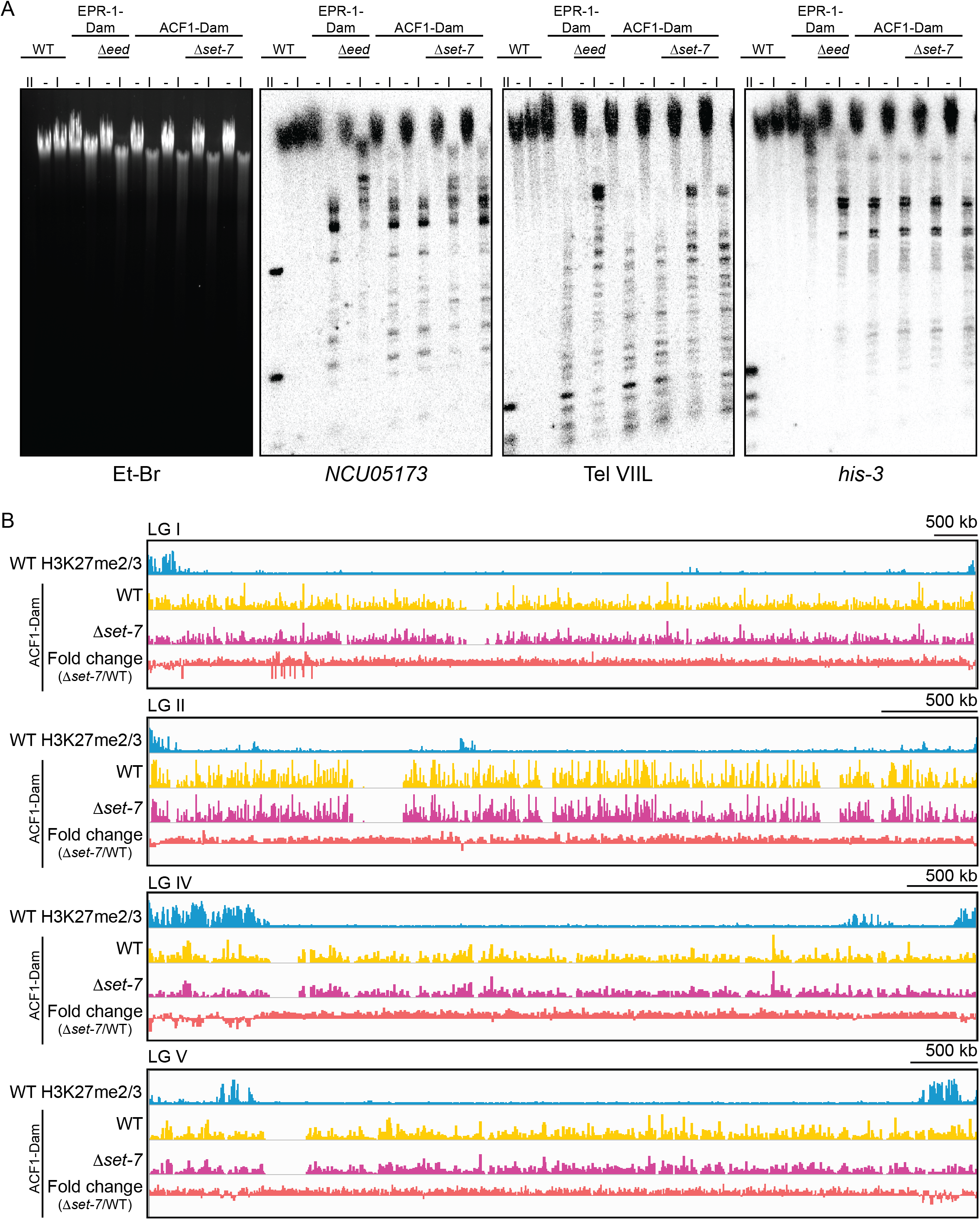
ACF1 localizes to H3K27me2/3 marked regions of the genome. A, DamID Southern blot with genomic DNA from the indicated strains digested with DpnI (I), DpnII (II) or left undigested (-). DpnII, which digests GATC sites without methylated adenines, shows the pattern of complete digestion in wild type. DpnI, which only digests GATC sites bearing adenine methylation, reveals the extent of methylation by Dam at probed regions (*NCU05173* and Tel VIIL, H3K27-methylated; *his-3*, euchromatin). Ethidium bromide (EtBr) shows total DNA. Biological replicates are shown for ACF1-Dam strains. EPR-1 is a presumptive H3K27 methyl-binding protein and was used as a positive control. EED and SET-7 are both members of the PRC2 complex and are required for H3K27 methylation. See source data for raw, uncropped images. B, Top track shows wild-type H3K27me2/3 levels averaged from two biological replicates of ChIP on the indicated linkage group. Y axis is 0-500 RPKM. Middle two tracks show DamID-seq average reads from two biological replicates for the indicated genotypes. Y axis is 0-500 RPKM. Bottom track compares the DamID-seq reads from Δ*set-7* to wild type (above) to show the fold change between the two genotypes. Y axis is −3 to 3.

**Figure 6 – figure supplement 1:**
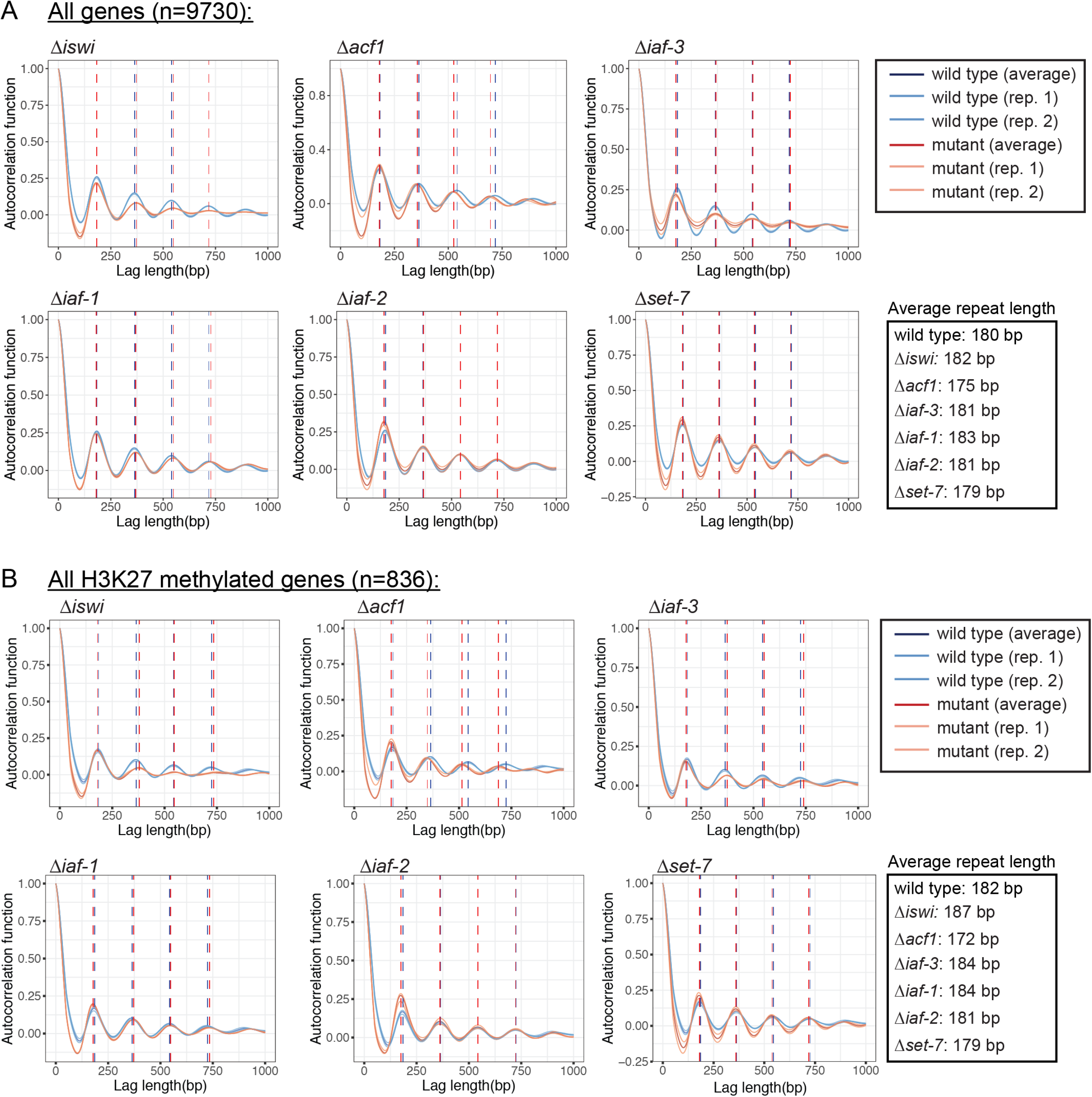
ISWI and its interacting partners have minor effects on nucleosome repeat length in *N. crassa*. A, Autocorrelation function is plotted for all genes (n=9730) in the indicated strains. Biological replicates and the average of the two replicates are shown. The vertical dotted blue line indicates wild-type nucleosome position and the vertical dotted red line indicates mutant nucleosome position. Average repeat length for each strain is shown on the right. B, Same as in (A), but for all H3K27 methylated genes (n=836).

**Figure 6 – figure supplement 2:**
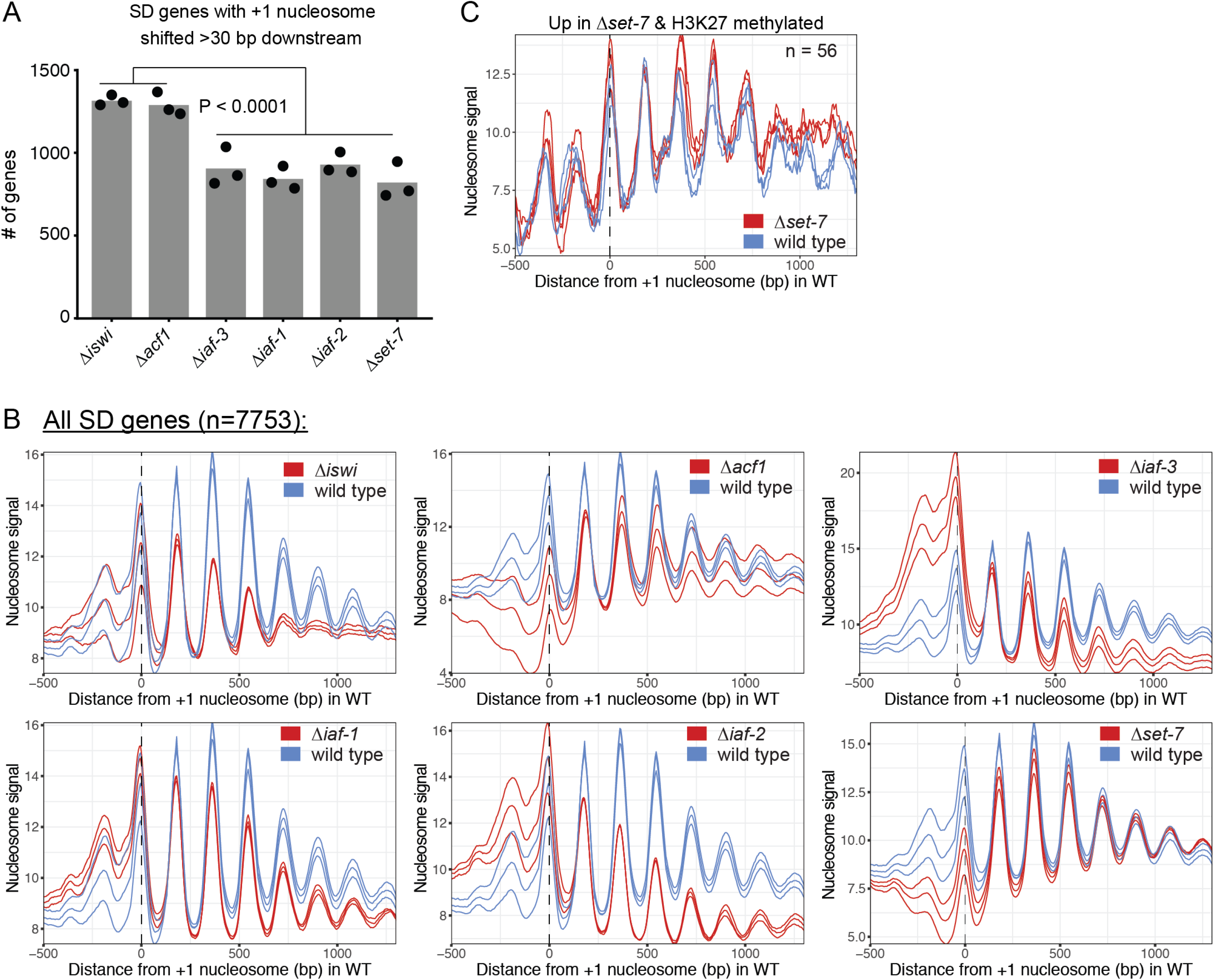
Global nucleosome positions in mutants and at genes upregulated in Δ*set-7*. A, Histogram of the number of SD genes (spectral density score for nucleosome order > 2; n=7753) that have the +1 nucleosome shifted downstream >30 base pairs when compared to wild type in the indicated mutant strains. Each point represents biological replicate 1, biological replicate 2, or analysis of the merged replicates and filled bar is the average of all three values. P values were determined with an unpaired t-test. B, Meta-analysis plot of the nucleosome signal at all SD genes. The three colored lines represent biological replicate 1, biological replicate 2, and the average of the replicates for the indicated strain. C, Average nucleosome signal at SD genes that are upregulated (FDR < 0.05) and marked by H3K27 methylation in Δ*set-7* strains. The three colored lines represent biological replicate 1, biological replicate 2, and the average of the replicates.

## SOURCE DATA LEGENDS

Figure 2 - source data 1: RNA-seq analysis (also applies to RNA-seq in Figure 3) Figure 3 – source data 1: ISWI interactor comparison total spectra greater than 0.4 from mass spectrometry

Figure 3 – source data 2: All mass spectrometry data

Figure 4 – source data 1: H3K27me2/3 ChIP-seq comparisons

Figure 5 – figure supplement 1 - source data 1: Raw image for Et-Br gel

Figure 5 – figure supplement 1 - source data 2: Raw image for Sothern blot probed with *NCU05173*

Figure 5 – figure supplement 1 - source data 3: Raw image for Southern blot probed with Tel VIIL

Figure 5 – figure supplement 1 - source data 4: Raw image for Southern blot probed with *his-3*

Figure 5 – figure supplement 1 - source data 5: Raw, uncropped image for Et-Br gel with labels

Figure 5 – figure supplement 1 - source data 6: Raw, uncropped image for Sothern blot probed with *NCU05173* with labels

Figure 5 – figure supplement 1 - source data 7: Raw, uncropped image for Sothern blot probed with Tel VIIL with labels

Figure 5 – figure supplement 1 - source data 8: Raw, uncropped image for Sothern blot probed with *his-3* with labels

Figure 6 – source data 1: List of SD genes used for MNase-seq analysis

---

**Supplementary File 1.**
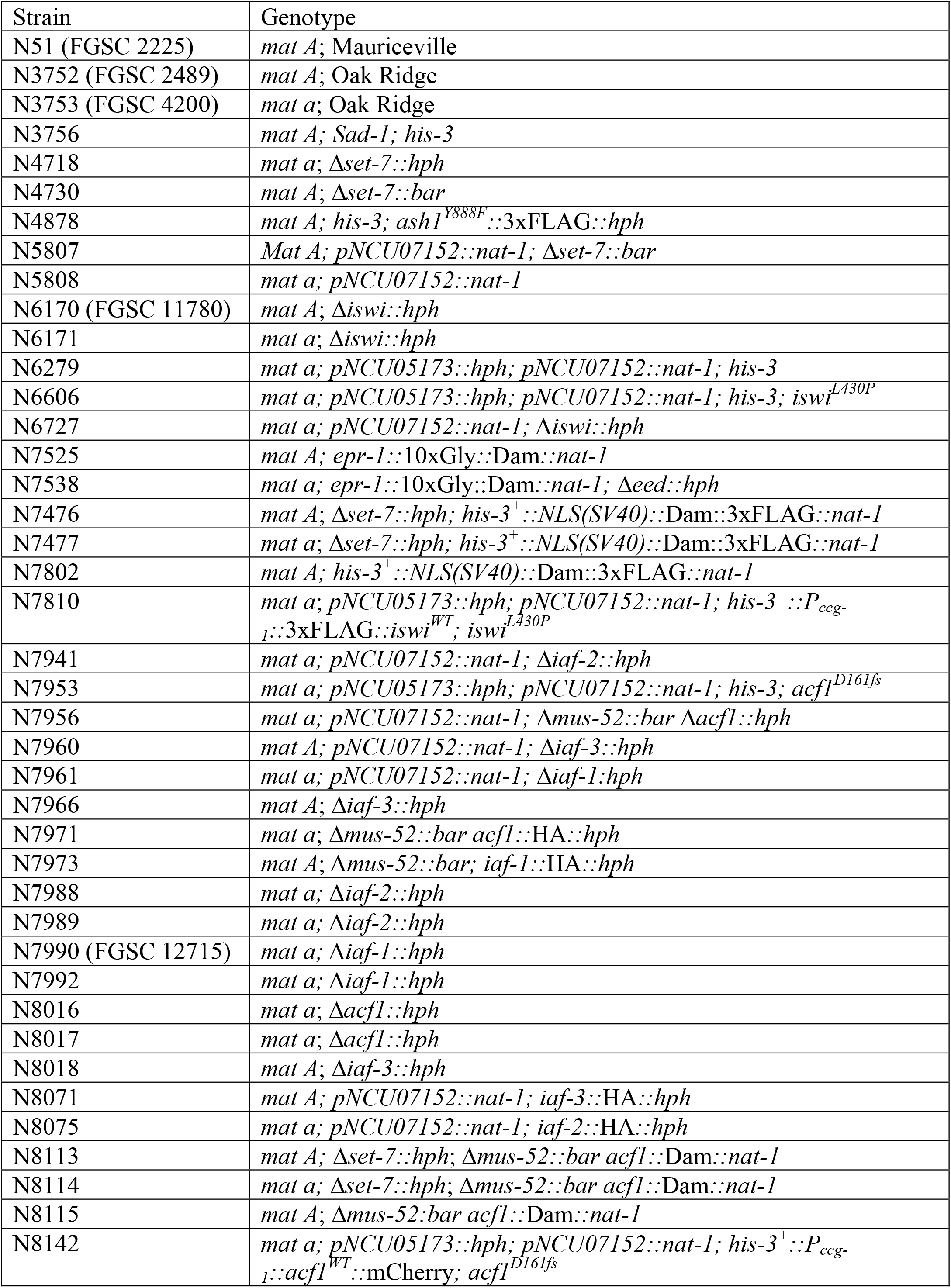
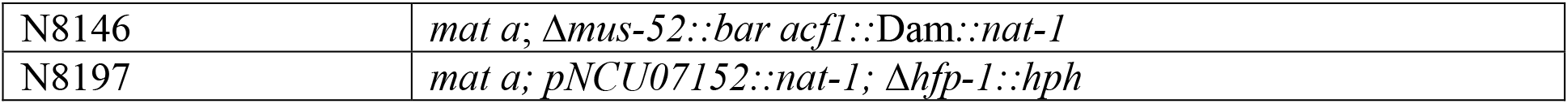
*Neurospora crassa* strains

**Supplementary File 2.**
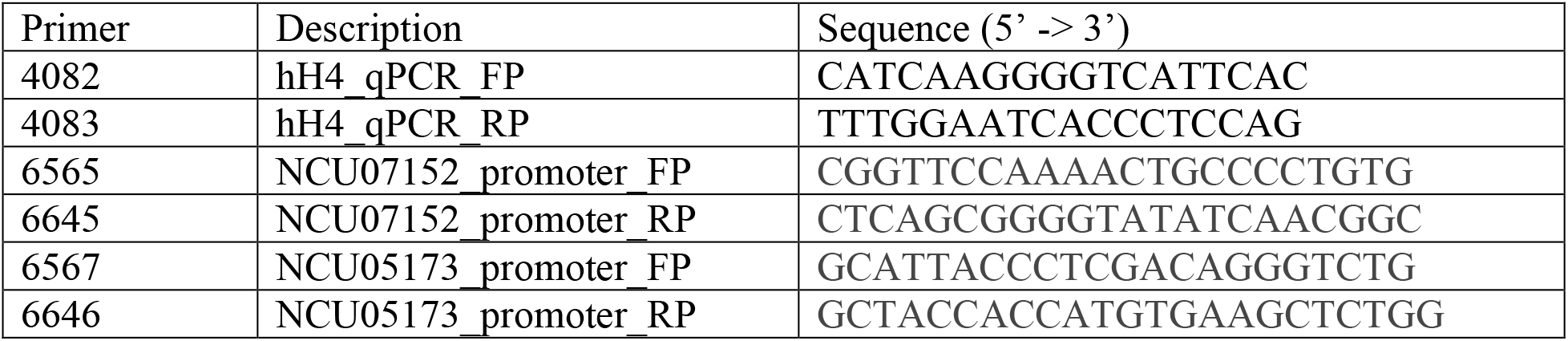
ChIP-qPCR primers

**Supplementary File 3.**
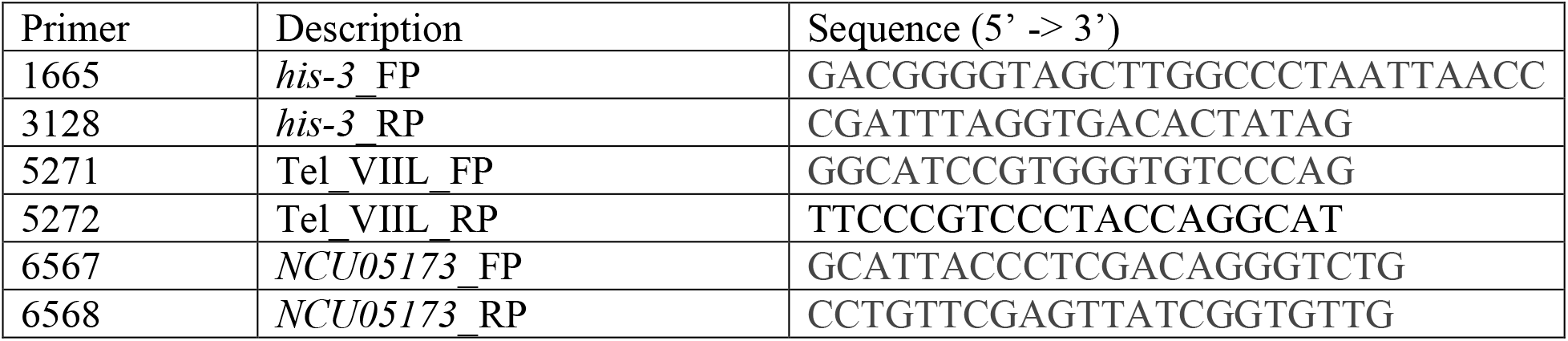
Southern probe primers

## REFERENCES

Afgan E, Baker D, Batut B, van den Beek M, Bouvier D, Cech M, Chilton J, Clements D, Coraor N, Gruning BA, et al. 2018. The Galaxy platform for accessible, reproducible and collaborative biomedical analyses: 2018 update. Nucleic Acids Res 46: W537–W544.

Altschul SF, Gish W, Miller W, Myers EW, Lipman DJ. 1990. Basic local alignment search tool. J Mol Biol 215: 403–410.

Baldi S, Jain DS, Harpprecht L, Zabel A, Scheibe M, Butter F, Straub T, Becker PB. 2018. Genome-wide Rules of Nucleosome Phasing in Drosophila. Mol Cell 72: 661–672 e664.

Bicocca VT, Ormsby T, Adhvaryu KK, Honda S, Selker EU. 2018. ASH1-catalyzed H3K36 methylation drives gene repression and marks H3K27me2/3-competent chromatin. Elife 7.

Borkovich KA, Alex LA, Yarden O, Freitag M, Turner GE, Read ND, Seiler S, Bell-Pedersen D, Paietta J, Plesofsky N et al. 2004. Lessons from the genome sequence of Neurospora crassa: tracing the path from genomic blueprint to multicellular organism. Microbiol Mol Biol Rev 68: 1–108.

Boyle S, Flyamer IM, Williamson I, Sengupta D, Bickmore WA, Illingworth RS. 2020. A central role for canonical PRC1 in shaping the 3D nuclear landscape. Genes Dev 34: 931–949.

Braunschweig U, Hogan GJ, Pagie L, van Steensel B. 2009. Histone H1 binding is inhibited by histone variant H3.3. EMBO J 28: 3635–3645.

Cheutin T, Cavalli G. 2018. Loss of PRC1 induces higher-order opening of Hox loci independently of transcription during Drosophila embryogenesis. Nat Commun 9: 3898.

Clapier CR, Cairns BR. 2012. Regulation of ISWI involves inhibitory modules antagonized by nucleosomal epitopes. Nature 492: 280–284.

Corona DF, Eberharter A, Budde A, Deuring R, Ferrari S, Varga-Weisz P, Wilm M, Tamkun J, Becker PB. 2000. Two histone fold proteins, CHRAC-14 and CHRAC-16, are developmentally regulated subunits of chromatin accessibility complex (CHRAC). EMBO J 19: 3049–3059.

Danecek P, Auton A, Abecasis G, Albers CA, Banks E, DePristo MA, Handsaker RE, Lunter G, Marth GT, Sherry ST et al. 2011. The variant call format and VCFtools. Bioinformatics 27: 2156–2158.

Donovan DA, Crandall JG, Truong VN, Vaaler AL, Bailey TB, Dinwiddie D, Banks OG, McKnight LE, McKnight JN. 2021. Basis of specificity for a conserved and promiscuous chromatin remodeling protein. Elife 10.

Erdel F, Schubert T, Marth C, Langst G, Rippe K. 2010. Human ISWI chromatin-remodeling complexes sample nucleosomes via transient binding reactions and become immobilized at active sites. Proc Natl Acad Sci U S A 107: 19873–19878.

Fazzio TG, Kooperberg C, Goldmark JP, Neal C, Basom R, Delrow J, Tsukiyama T. 2001. Widespread collaboration of Isw2 and Sin3-Rpd3 chromatin remodeling complexes in transcriptional repression. Mol Cell Biol 21: 6450–6460.

Fyodorov DV, Blower MD, Karpen GH, Kadonaga JT. 2004. Acf1 confers unique activities to ACF/CHRAC and promotes the formation rather than disruption of chromatin in vivo. Genes Dev 18: 170–183.

Garrison E, Marth G. 2012. Haplotype-based variant detection from short-read sequencing. arXiv.

Gelbart ME, Bachman N, Delrow J, Boeke JD, Tsukiyama T. 2005. Genome-wide identification of Isw2 chromatin-remodeling targets by localization of a catalytically inactive mutant. Genes Dev 19: 942–954.

Goldmark JP, Fazzio TG, Estep PW, Church GM, Tsukiyama T. 2000. The Isw2 chromatin remodeling complex represses early meiotic genes upon recruitment by Ume6p. Cell 103: 423–433.

Grau DJ, Chapman BA, Garlick JD, Borowsky M, Francis NJ, Kingston RE. 2011. Compaction of chromatin by diverse Polycomb group proteins requires localized regions of high charge. Genes Dev 25: 2210–2221.

Hunter JD. 2007. Matplotlib: A 2D Graphics Environment. Computing in Science & Engineering 9: 90–95.

Hwang WL, Deindl S, Harada BT, Zhuang X. 2014. Histone H4 tail mediates allosteric regulation of nucleosome remodelling by linker DNA. Nature 512: 213–217.

Iida T, Araki H. 2004. Noncompetitive counteractions of DNA polymerase epsilon and ISW2/yCHRAC for epigenetic inheritance of telomere position effect in Saccharomyces cerevisiae. Mol Cell Biol 24: 217–227.

Ito T, Bulger M, Pazin MJ, Kobayashi R, Kadonaga JT. 1997. ACF, an ISWI-containing and ATP-utilizing chromatin assembly and remodeling factor. Cell 90: 145–155.

Ito T, Levenstein ME, Fyodorov DV, Kutach AK, Kobayashi R, Kadonaga JT. 1999. ACF consists of two subunits, Acf1 and ISWI, that function cooperatively in the ATP-dependent catalysis of chromatin assembly. Genes Dev 13: 1529–1539.

Jamieson K, McNaught KJ, Ormsby T, Leggett NA, Honda S, Selker EU. 2018. Telomere repeats induce domains of H3K27 methylation in Neurospora. Elife 7.

Jamieson K, Rountree MR, Lewis ZA, Stajich JE, Selker EU. 2013. Regional control of histone H3 lysine 27 methylation in Neurospora. Proc Natl Acad Sci U S A 110: 6027–6032.

Kamei M, Ameri AJ, Ferraro AR, Bar-Peled Y, Zhao F, Ethridge CL, Lail K, Amirebrahimi M, Lipzen A, Ng V et al. 2021. IMITATION SWITCH is required for normal chromatin structure and gene repression in PRC2 target domains. Proc Natl Acad Sci U S A 118.

Kassis JA, Brown JL. 2013. Polycomb group response elements in Drosophila and vertebrates. Adv Genet 81: 83–118.

Kassis JA, Kennison JA, Tamkun JW. 2017. Polycomb and Trithorax Group Genes in Drosophila. Genetics 206: 1699–1725.

Klocko AD, Ormsby T, Galazka JM, Leggett NA, Uesaka M, Honda S, Freitag M, Selker EU. 2016. Normal chromosome conformation depends on subtelomeric facultative heterochromatin in Neurospora crassa. Proc Natl Acad Sci U S A 113: 15048–15053.

Kornberg RD. 1974. Chromatin structure: a repeating unit of histones and DNA. Science 184: 868–871.

Kubik S, Bruzzone MJ, Challal D, Dreos R, Mattarocci S, Bucher P, Libri D, Shore D. 2019. Opposing chromatin remodelers control transcription initiation frequency and start site selection. Nat Struct Mol Biol 26: 744–754.

Kubik S, O’Duibhir E, de Jonge WJ, Mattarocci S, Albert B, Falcone JL, Bruzzone MJ, Holstege FCP, Shore D. 2018. Sequence-Directed Action of RSC Remodeler and General Regulatory Factors Modulates +1 Nucleosome Position to Facilitate Transcription. Mol Cell 71: 89–102 e105.

Lai WKM, Pugh BF. 2017. Understanding nucleosome dynamics and their links to gene expression and DNA replication. Nat Rev Mol Cell Biol 18: 548–562.

Langmead B, Salzberg SL. 2012. Fast gapped-read alignment with Bowtie 2. Nat Methods 9: 357–359.

Laura E McKnight JGC, Thomas B Bailey, Orion GB Banks, Kona N Orlandi, Vi N Truong, Grace L Waddell, Elizabeth T Wiles, Drake A Donovan, Scott D Hansen, Eric U Selker, Jeffrey N McKnight. 2019. Rapid and Inexpensive Preparation of Genome-Wide Nucleosome Footprints from Model and Non-Model Organisms. BioRxiv 870659.

LeRoy G, Loyola A, Lane WS, Reinberg D. 2000. Purification and characterization of a human factor that assembles and remodels chromatin. J Biol Chem 275: 14787–14790.

Ludwigsen J, Pfennig S, Singh AK, Schindler C, Harrer N, Forne I, Zacharias M, Mueller-Planitz F. 2017. Concerted regulation of ISWI by an autoinhibitory domain and the H4 N-terminal tail. Elife 6.

Luger K, Mader AW, Richmond RK, Sargent DF, Richmond TJ. 1997. Crystal structure of the nucleosome core particle at 2.8 A resolution. Nature 389: 251–260.

Margueron R, Reinberg D. 2011. The Polycomb complex PRC2 and its mark in life. Nature 469: 343–349.

McNaught KJ, Wiles ET, Selker EU. 2020. Identification of a PRC2 Accessory Subunit Required for Subtelomeric H3K27 Methylation in Neurospora crassa. Mol Cell Biol 40.

Metzenberg RL, Stevens JN, Selker EU, Morzycka-Wroblewska E. 1985. Identification and chromosomal distribution of 5S rRNA genes in Neurospora crassa. Proc Natl Acad Sci U S A 82: 2067–2071.

Miao VP, Freitag M, Selker EU. 2000. Short TpA-rich segments of the zeta-eta region induce DNA methylation in Neurospora crassa. J Mol Biol 300: 249–273.

Muller J, Hart CM, Francis NJ, Vargas ML, Sengupta A, Wild B, Miller EL, O’Connor MB, Kingston RE, Simon JA. 2002. Histone methyltransferase activity of a Drosophila polycomb group repressor complex. Cell 111: 197–208.

Nishioka K, Miyazaki H, Soejima H. 2018. Unbiased shRNA screening, using a combination of FACS and high-throughput sequencing, enables identification of novel modifiers of Polycomb silencing. Sci Rep 8: 12128.

Nocetti N, Whitehouse I. 2016. Nucleosome repositioning underlies dynamic gene expression. Genes Dev 30: 660–672.

Ocampo J, Chereji RV, Eriksson PR, Clark DJ. 2016. The ISW1 and CHD1 ATP-dependent chromatin remodelers compete to set nucleosome spacing in vivo. Nucleic Acids Res 44: 4625–4635.

Petty E, Pillus L. 2013. Balancing chromatin remodeling and histone modifications in transcription. Trends Genet 29: 621–629.

Pomraning KR, Smith KM, Freitag M. 2011. Bulk segregant analysis followed by high-throughput sequencing reveals the Neurospora cell cycle gene, ndc-1, to be allelic with the gene for ornithine decarboxylase, spe-1. Eukaryot Cell 10: 724–733.

Rhee HS, Pugh BF. 2012. Genome-wide structure and organization of eukaryotic pre-initiation complexes. Nature 483: 295–301.

Ridenour JB, Moller M, Freitag M. 2020. Polycomb Repression without Bristles: Facultative Heterochromatin and Genome Stability in Fungi. Genes (Basel) 11.

Scacchetti A, Brueckner L, Jain D, Schauer T, Zhang X, Schnorrer F, van Steensel B, Straub T, Becker PB. 2018. CHRAC/ACF contribute to the repressive ground state of chromatin. Life Sci Alliance 1: e201800024.

Schuettengruber B, Bourbon HM, Di Croce L, Cavalli G. 2017. Genome Regulation by Polycomb and Trithorax: 70 Years and Counting. Cell 171: 34–57.

Tsukiyama T, Palmer J, Landel CC, Shiloach J, Wu C. 1999. Characterization of the imitation switch subfamily of ATP-dependent chromatin-remodeling factors in Saccharomyces cerevisiae. Genes Dev 13: 686–697.

van Steensel B, Henikoff S. 2000. Identification of in vivo DNA targets of chromatin proteins using tethered dam methyltransferase. Nat Biotechnol 18: 424–428.

Varga-Weisz PD, Wilm M, Bonte E, Dumas K, Mann M, Becker PB. 1997. Chromatin-remodelling factor CHRAC contains the ATPases ISWI and topoisomerase II. Nature 388: 598–602.

Vary JC, Jr., Gangaraju VK, Qin J, Landel CC, Kooperberg C, Bartholomew B, Tsukiyama T. 2003. Yeast Isw1p forms two separable complexes in vivo. Mol Cell Biol 23: 80–91.

Weber CM, Ramachandran S, Henikoff S. 2014. Nucleosomes are context-specific, H2A.Z-modulated barriers to RNA polymerase. Mol Cell 53: 819–830.

Whitehouse I, Rando OJ, Delrow J, Tsukiyama T. 2007. Chromatin remodelling at promoters suppresses antisense transcription. Nature 450: 1031–1035.

Whitehouse I, Tsukiyama T. 2006. Antagonistic forces that position nucleosomes in vivo. Nat Struct Mol Biol 13: 633–640.

Wiles ET, McNaught KJ, Kaur G, Selker JML, Ormsby T, Aravind L, Selker EU. 2020. Evolutionarily ancient BAH-PHD protein mediates Polycomb silencing. Proc Natl Acad Sci U S A 117: 11614–11623.

Wiles ET, Selker EU. 2017. H3K27 methylation: a promiscuous repressive chromatin mark. Curr Opin Genet Dev 43: 31–37.

Yen K, Vinayachandran V, Batta K, Koerber RT, Pugh BF. 2012. Genome-wide nucleosome specificity and directionality of chromatin remodelers. Cell 149: 1461–1473.

Zhou V. 2012. Methods for Global Characterization of Chromatin Regulators in Human Cells. Harvard University Press.

